# H_2_S mediates interbacterial communication through the air reverting intrinsic antibiotic resistance

**DOI:** 10.1101/202804

**Authors:** Daniel Thomas-Lopez, Laura Carrilero, Stephanie Matrat, Natalia Montero, Stéphane Claverol, Milos R Filipovic, Bruno Gonzalez-Zorn

## Abstract

Hydrogen sulfide, a gas classically considered as a by-product of cellular metabolism, is today recognized as a crucial gasotransmitter in Eukaryotes. Moreover, most bacteria harbor the eukaryotic orthologous genes for H_2_S synthesis, and these genes have been linked to different metabolic pathways.

Some bacteria, however, produce high amounts of H_2_S in their extracellular space, a characteristic classically used for identification purposes. This is the case of *Salmonella* Typhimurium, which produces H_2_S by its *phsABC* operon. Here we show that extracellular release of H_2_S by *S*. Typhimurium is solely dependent on its *phsABC* operon. Furthermore, we show that *S*. Typhimurium and other H_2_S-producing bacteria can interact with physically distant bacteria through H_2_S production. We demonstrate how H_2_S can revert intrinsic cephalosporin resistance of *Enterococccus faecalis* and *Enterococcus faecium* to complete susceptibility. This study constitutes a significant step in the study of bacterial interplay and niche competition. Furthermore, as H_2_S releasing drugs have already been designed, our results open the way to future therapeutic alternatives for the treatment of infections caused by enterococci, multiresistant pathogens for which no treatments are clinically available.

**Author Summary:** It has been known for decades that bacteria can communicate with each other through the diffusion of metabolites in the media. However, the capacity of a bacterium to interact with other physically distant cell is a recent discovery of the 21^st^ century. In this work we show how some well-studied bacteria, as it is *Salmonella* spp., interacts with other bacteria thanks to the compound hydrogen sulfide (H_2_S) that they produce and release to the environment.

In our study we have designed novel techniques that allow us to study the interaction between two bacteria, and we have seen that *Salmonella* is able to affect other species that is even 1 cm away, *i.e.*, a distance corresponding to 10.0000 times its own size.

What is more astonishing is that *Enterococcus*, when exposed to the H_2_S, is dramatically becomes susceptible to many antibiotics, to which it is supposed to be naturally resistant. *Enterococcus* spp. are responsible for life-threatening infections in hospitals worldwide. Thus, our observations reveal that bacteria can communicate through the air with H2S, and that this molecule can make bacteria that are highly resistant to antibiotics susceptible to antibiotics, making untreatable infections treatable with current antibiotics.

## Introduction

It is well established that bacteria can communicate with each other through the diffusion of molecules in the media, what we know as quorum sensing [1]. On the other hand, in the 21^st^ century it has been discovered that physically distant bacteria can also interact with each other or with distant organisms by releasing gaseous molecules [2], many of which were previously considered mere byproducts of bacterial metabolism. Ammonia [3], indole [4], trimethylamine [4] or acetic acid [5] are some these volatile compounds responsible for changes in motility, biofilm formation or antibiotic resistance. In addition, there are a few gases, called gasotransmitters, that are particularly interesting as they also play important roles in eukaryotic cells [6]. Gasotransmitters include mainly nitric oxide (NO), hydrogen sulfide (H_2_S) and carbon monoxide (CO), and they have been connected to physiological and pathological conditions in cancer [7], the cardiovascular system [8,9], potassium channels [10], cellular ageing [11], animal hibernation [12] and grapes’ senescence [13], among others.

In microbiology, NO and H_2_S have received special attention as they enhance global antimicrobial resistance [14,15]. Bacteria have been shown to possess the orthologous genes to those found in eukaryotic cells for the production of these gases.

In the case of hydrogen sulfide, Shatalin *et al.* revealed that this gas confers intracellularly general protection against the bactericidal action of antibiotics through the eukaryotic orthologous pathways cystathionine-β-synthase (CBS), cystathionine-γ-lyase (CSE) or 3-mercaptopyruvate sulfurtransferase (3MST), present in many bacterial families of clinical interest [15].

However, H_2_S production in large amounts by specific pathways found in certain bacteria, *e*. *g*. from the *Salmonella, Citrobacter, Edwardsiella* and *Proteus* genera, has been well known for more than 50 years [16]. Of the different H_2_S synthesis mechanisms in bacteria, the most accurately characterized is the *phsABC* operon of *Salmonella* Typhimurium, which generates H_2_S and sulfite through thiosulfate reduction. The purpose of this H_2_S synthesis by *S*. Typhimurium is not well understood. It is well established that in the host’s gut, thiosulfate can be oxidized to tetrathionate, which can be used by *S*. Typhimurium for respiratory purposes (reducing it again to thiosulfate) thanks to the *ttr* genes located in the pathogenicity island 2 (SPI-2) [17]. However, the benefits of a further reduction of thiosulfate to H_2_S (instead of using it for further tetrathionate production) are very scarce [18].

Besides, if the function of this gas in bacteria is to enhance antibiotic resistance [15,19,20], it is intriguing that some species, like the gram positive *Enterococcus*, inhibit H_2_S production by others [21]. However, this suggests that H_2_S might be interacting with neighboring microorganisms, as it is the case with other molecules already mentioned.

Here we characterize the implication of the *phsABC* operon in H_2_S synthesis by *S*. Typhimurium. As H_2_S produced by the PHS pathway is released extracellularly, we studied if *S*. Typhimurium could communicate with other bacteria thanks to its H_2_S production. We observed that it is not only the interaction that takes place but that *S*. Typhimurium can also revert other bacteria intrinsic antibiotic resistance, suggesting niche competition situations can be more complex than expected. Finally, we also demonstrate that H_2_S abolishes antibiotic resistance. As H_2_S releasing donors have already been developed, the combined application of antibiotics with H_2_S requires future in depth studying.

## Results

### H_2_S excretion relies entirely on *phsABC* action

Most bacteria possess H_2_S producing genes, namely CBS/CSE and 3MST [15], together with cysteine desulfhydrases [22]. Nevertheless, a few bacterial species carry, in addition to the previously mentioned genes, an accessory and specific H_2_S producing pathway. For example, *Salmonella* Typhimurium harbors the *phsABC* operon. To understand the role of this pathway, we deleted this operon (Fig 1A) and we measured the H_2_S production of the deletion mutant *Δphs*, both in aerobic and anaerobic conditions.

**Fig 1.**
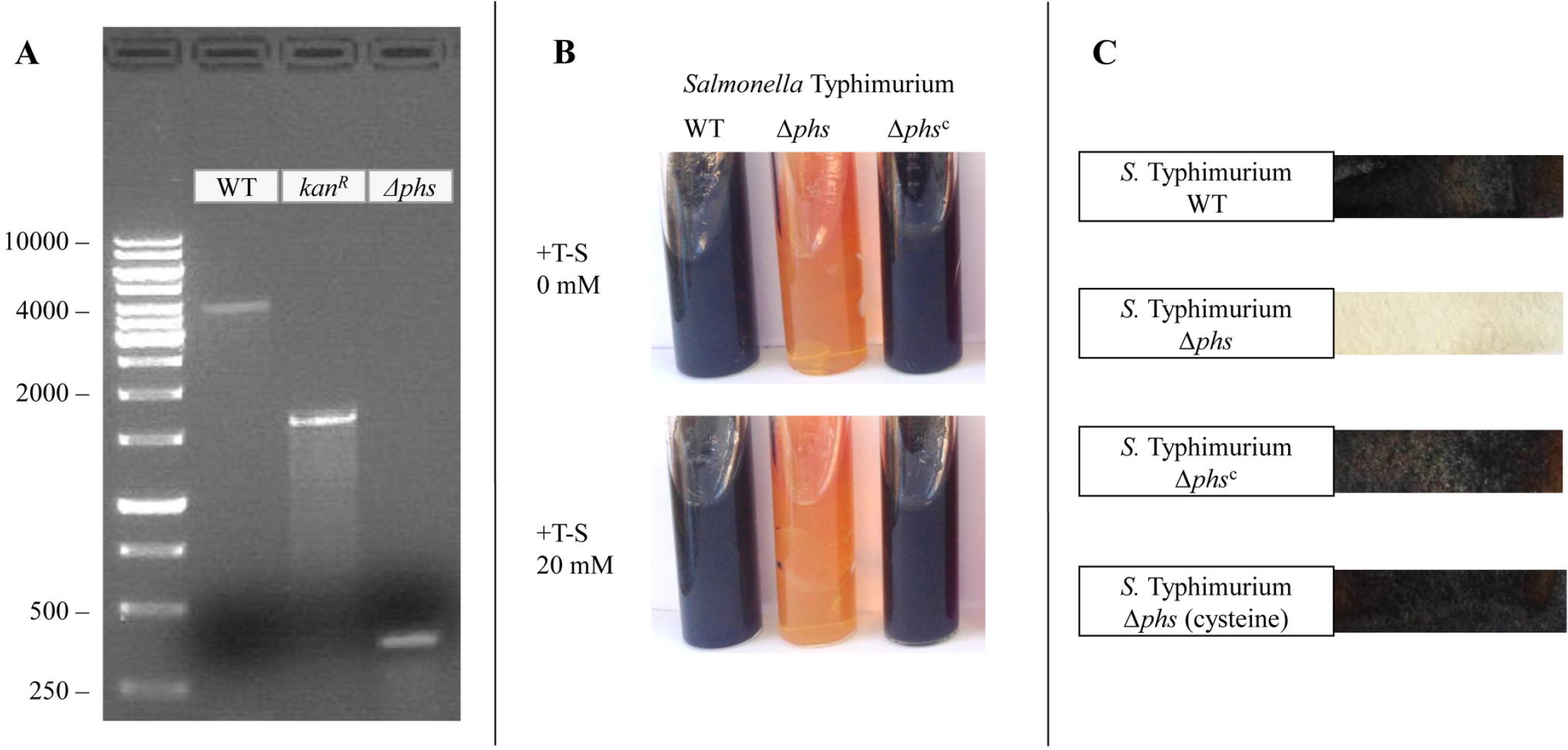
Characterization of the *S.* Typhimurium *phsABC* operon. (A) PCR showing substitution of the operon by the kanamycin cassette, and subsequent deletion of this cassette. (B) H_2_S production, measured in Kligler media in the absence (upper line) or the presence (bottom line) of thiosulfate 20 mM (C) or in peptone water broth by lead acetate strips.

Kligler media, which in the presence of H_2_S forms a dark precipitate, turned completely black when *S*. Typhimurium wild-type strain was inoculated, whereas *Δphs* did not produce any black pigmentation of the media at all (Fig 1B).

Lead (II) acetate paper strips, which detect H_2_S through the formation of black lead sulfide, allowed us to detect H_2_S released not only to the media, but also outside of the media, as the strips were located above the culture medium. Again, we observed that the strips turned completely black with the WT culture, while incubation of *Δphs* strain barely caused any staining (Fig 1C).

In both cases, complementation of the strain with the pSB74 plasmid, which will be called pH_2_S as it bears the entire *phsABC* operon [23], restored H_2_S production at WT levels in the new strain *S*. Typhimurium *Δphs*^c^ (Fig 1).

### The PHS pathway is neither involved in growth nor antibiotic resistance in *S*. Typhimurium

H_2_S has been linked to antibiotic resistance, via the H_2_S synthesis pathways found in most bacteria [15,20]. However, this has not been demonstrated for the *S*. Typhimurium specific PHS pathway, even though this is considered the main source of H_2_S in this species [23,24]. Therefore, we carried out various antimicrobial susceptibility tests (S1 Fig, **Error! Reference source not found.**S1 and S2 Tables), but no differences were observed in any case. Growth curves showed no difference either in growth rate between WT and *Δphs* strain (S1 Fig).

### Remote action of *S.* Typhimurium on the intrinsic antibiotic resistance of *Enterococcus*

H_2_S produced by means of the PHS pathway by *S*. Typhimurium is largely released from the cell and does not appear to be implicated its own growth or antibiotic resistance. Thus, we hypothesized that H_2_S may have an external role on the antibiotic resistance pattern of neighboring bacteria when *S*. Typhimurium interacted with them.

We designed a model to effectively assess signaling between physically distant bacteria (Fig 2). This method consists in preparing antibiograms of the species to be tested and confronting them against another Petri dish in which the H_2_S producing strain has been plated with a cotton swab. Subsequently, plates were incubated for a maximum of 24 hours.

**Fig 2.**
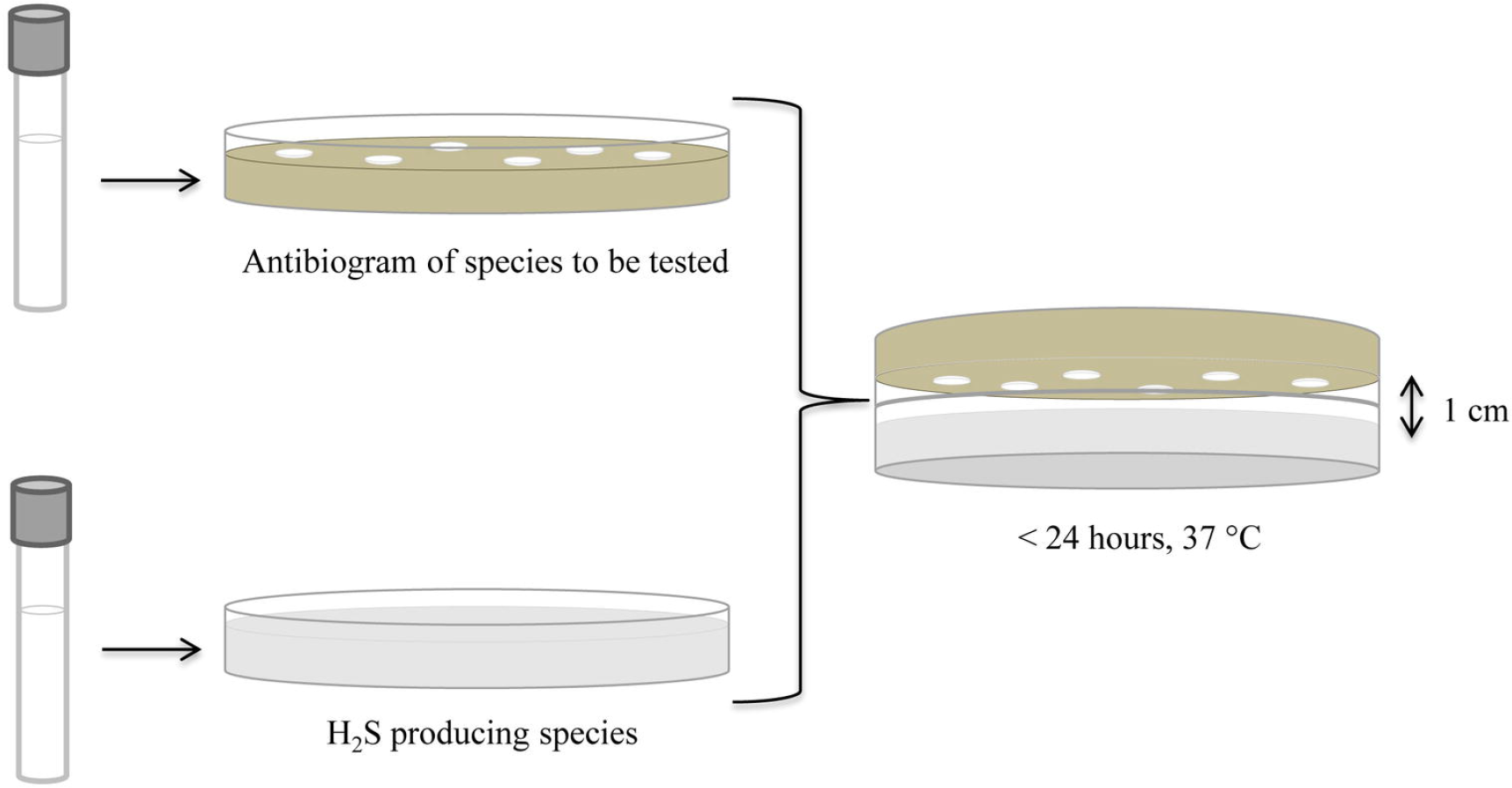
Confronting experiments. An antibiogram of the species to be tested is exposed to another plate streaked with an H_2_S producing species and supplemented with thiosulfate 20 mM (T-S).

*S*. Typhimurium did not significantly alter bacterial growth, hemolysis capacity, colonies’ size and morphology or the antibiotic sensitivity profile of the species analyzed, which includes methicillin-resistant *Staphylococcus aureus, Bacillus cereus, Escherichia coli, Proteus vulgaris* and the *S*. Typhimurium *Δphs* mutant itself (data not shown).

However, when faced with *S*. Typhimurium, *E. faecalis*, a major nosocomial pathogen worldwide, became completely susceptible against cephalosporins, drugs to which *Enterococcus* is intrinsically resistant (Fig 3]).

**Fig 3.**
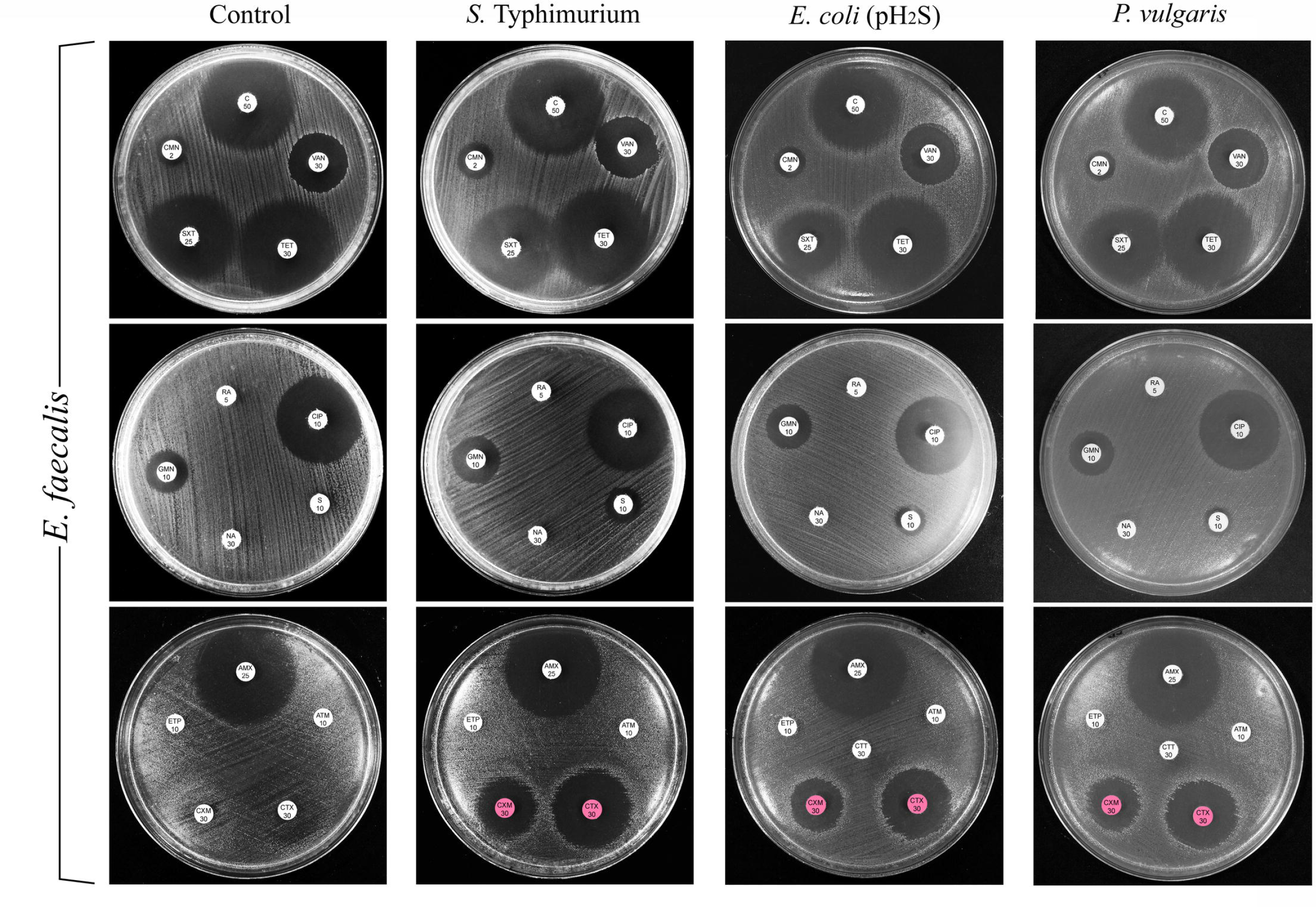
Confronting experiments of *E*. *faecalis*. Control antibiograms and confronting experiments of *E*. *faecalis* JH2-2 (plated on TSA media) faced with *S*. Typhimurium ATCC14028, *E*. *coli* (pH_2_S), *Proteus vulgaris* CECT174. H_2_S producing species were plated on TSA media containing T-S 20 mM. Antibiotics tested: cloramphenicol (C), vancomycin (VAN), tetracycline (TET), sulphametoxazol-trimethoprim (SXT), clindamycin (CMN), rifampin (RA), ciprofloxacin (CIP), streptomycin (S), nalidixic acid (NA), gentamicin (GMN), amoxicillin (AMX), aztreonam (ATM), cefotaxime (CTX), cefuroxime (CXM), ertapenem (ETP), cefotetan (CTT). Antibiotics showing synergy with H_2_S are highlighted.

### The *phsABC* operon of *Salmonella* is responsible for the reversion of *Enterococcus* intrinsic antibiotic resistance

To distinguish if enterococci lose their cephalosporin resistance due to H_2_S released by *S*. Typhimurium or by an unrelated mechanism, we confronted *E. faecalis* with the deletion mutant *Δphs*, which we have shown not to release any H_2_S.

By performing E-test, we observed that *E. faecalis* Minimal Inhibitory Concentration (MIC) for cefotaxime was dramatically decreased, from higher than 256 mg/L to 3 mg/L, when faced specifically with *S*. Typhimurium WT, *i*. *e*., the H_2_S producing strain. On the other hand, the MIC was not affected when faced with *S*. Typhimurium *Δphs* strain (Fig 4).

**Fig 4.**
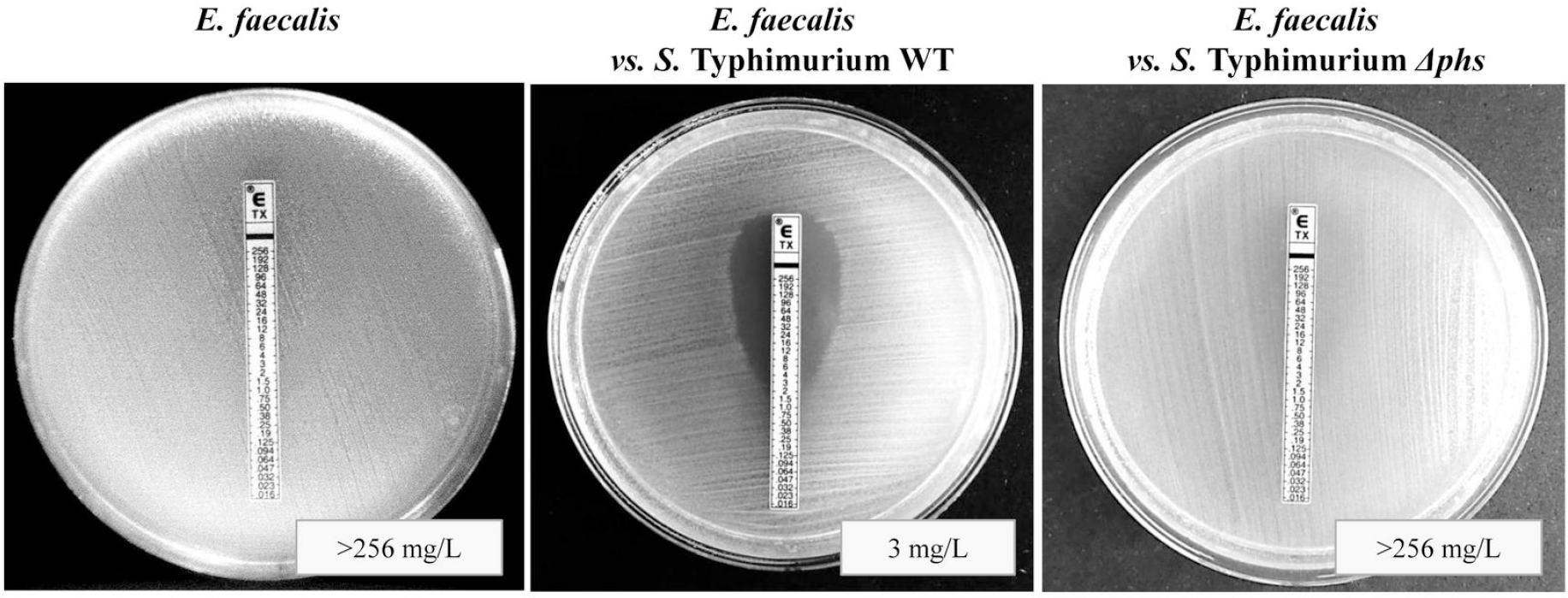
Minimal Inhibitory Concentration of *E*. *faecalis* confronted with *S.* Typhimurium. MIC for cefotaxime (TX) of *E*. *faecalis* JH2-2 control strain (left), confronted with *S*. Typhimurium WT (center) and with *S*. Typhimurium *Δphs* (right).

*Δphs* mutant recovered its lethal action against *E. faecalis* when was complemented *in trans* (Table 1), which means that it is specifically via the H_2_S produced by the *phsABC* operon that *S*. Typhimurium induces *Enterococcus* killing. *S*. Typhimurium (pTrc99a), *i*.*e*., the strain carrying the empty vector pTrc99a, was used as a negative control to ensure that this plasmid had no influence on the effect. As we established that *phsABC* is a predominant mechanism for extracellular H_2_S release from this species, we conclude that this operon is crucial for the remote induced killing of neighbor bacteria by *S*. Typhimurium.

**Table 1.** **Confronting experiments of *E. faecalis* with *S.* Typhimurium WT and its respective mutants.**

| Strains | + T-S 2 mM |  |  | + T-S 20 mM |  |  | + Cys 2 mM |  |  | + T-S 20 mM Anaerobiosis |  |  |
| --- | --- | --- | --- | --- | --- | --- | --- | --- | --- | --- | --- | --- |
|  | ZOX | CTX | CEC |  |  |  |  |  |  |  |  |  |
| *E. faecalis* | 6 | 6 | 16 |  |  | - |  |  | - |  |  | - |
| *vs. S. Typhimurium* WT | <u>28</u> | <u>30</u> | 19 | <u>25</u> | <u>33</u> | 16 | <u>27</u> | <u>30</u> | 17 | <u>22</u> | <u>26</u> | 19 |
| *vs. S. Typhimurium* $\Delta phs$ | 6 | 6 | 14 | 6 | 6 | 12 | <u>27</u> | <u>30</u> | 17 | 6 | 6 | 14 |
| *vs. $\Delta phs$ (pTrc99a)* | 6 | 6 | 14 | 6 | 6 | 13 | <u>23</u> | <u>23</u> | 16 |  |  | ND |
| *vs. $\Delta phs$ (pH<sub>2</sub>S)* | <u>25</u> | <u>28</u> | 16 | <u>32</u> | <u>34</u> | 16 | <u>25</u> | <u>27</u> | 17 |  |  | ND |

ND: not determined

*E. faecalis* JH2-2 is plated on Tryptone-Soya Agar (TSA). *S*. Typhimurium strains are plated on TSA supplemented with thiosulfate 2 or 20 mM. Halos are expressed in mm. Inhibition zones displayed only in the presence of H_2_S are underlined. Ceftizoxime (ZOX), Cefotaxime (CTX), Cefaclor (CEC).

### The *phsABC* operon is sufficient to induce cephalosporin susceptibility in *E. faecalis*

To identify if the product of the *phsABC* operon was having an effect in adjuvancy with other factors from *S*. Typhimurium, we transformed *E*. *coli* MG1655 strain with the pH_2_S plasmid. We show that the *phsABC* operon is sufficient for the increased production of H_2_S by *E*. *coli* MG1655 (Fig 5), implying that no further networks or genes present in *Salmonella* are necessary. Lead (II) acetate paper strips were stained in a similar way in *E*. *coli* (pH_2_S) and *S*. Typhimurium. When *E*. *coli* (pH_2_S) was incubated in Kligler media, we observed a lesser amount of staining than in *S*. Typhimurium. This can be explained by the fact that *S*. Typhimurium also harbors the anaerobic sulfite reductase (*asr*) that generates further H_2_S from the reduction of the sulfite previously generated by PHS.

**Fig 5.**
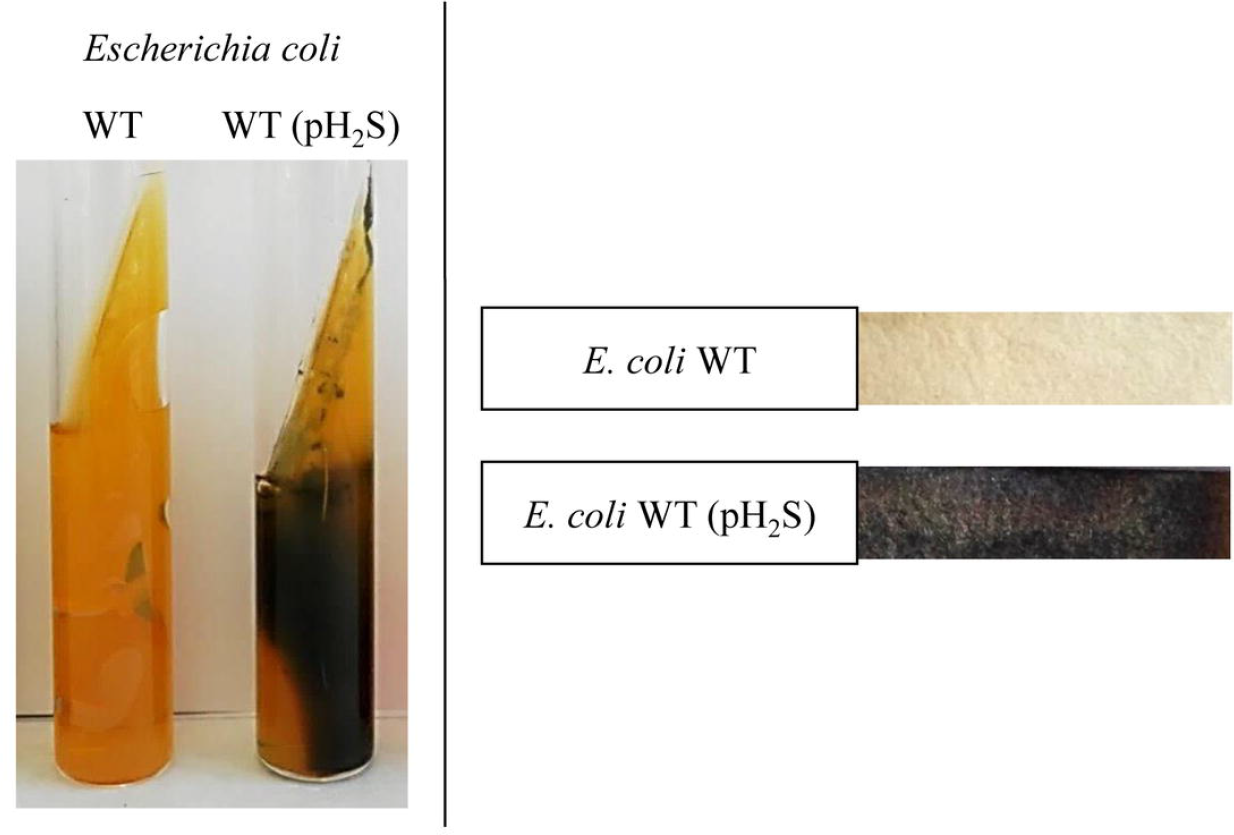
H_2_S production by *E*. *coli* WT and *E*. *coli* p(H_2_S). H_2_S production was measured with Kligler media (left) and lead acetate strips (right).

Furthermore, when we confronted different *E*. *coli* strains with *E. faecalis*, we observed that only *E*. *coli* (pH_2_S) could induce the susceptibility of *E. faecalis* to cephalosporins (Fig 3), showing that the *phsABC* function is sufficient for the resistance reversion phenomenon carried out by H_2_S and that no secondary metabolites produced by *S*. Typhimurium are necessary.

We also tested if our observations could be dependent on the presence of oxygen. However, the confrontations experiments performed in aerobiosis and anaerobiosis showed similar results (Table 1), discarding therefore that our results are connected to an augmentation of the reactive oxidative species.

### H_2_S is responsible for the reversion of the intrinsic resistance to cephalosporins in *E. faecalis*

The deletion of the *phsABC* operon of *S*. Typhimurium could have further consequences on the bacteria (*e*. *g*., affecting other metabolic pathways) that could ultimately be responsible for the distant effect observed on *Enterococcus* rather than the H_2_S itself. To rule out this hypothesis, we tested other sources of H_2_S. First, we carried out confrontation experiments in the presence of cysteine, the substrate of the 3MST pathway [15], present in many bacteria including *S*. Typhimurium. Even if we have proved that under basal conditions PHS is the predominant mechanism for extracellular production of H_2_S, we hypothesized that addition of external cysteine should increase the production of H_2_S by this pathway, as high levels of cysteine are toxic for the cell [22]. Upon adding cysteine, we observed that H_2_S is again detected extracellularly in the *Δphs* mutant (Fig 1) and that this mutant is able to revert *Enterococcus* cephalosporin resistance in a similar extent as the wild-type *S*. Typhimurium (Table 1). We confronted *E*. *faecalis* with *Proteus vulgaris*, another species commonly known for its extracellular production of H_2_S [22] and *P*. *vulgaris* proved to be equally efficient in inducing cephalosporin susceptibility in *E*. *faecalis* (Fig 3).

Finally, we checked if pure H_2_S from a chemical origin also acts synergistically with cephalosporins. NaHS is a widely used H_2_S source [15,25]. We designed a technique in which we placed NaHS physically separated from the enterococci. We put our antibiograms in a glass chamber, next to an empty Petri dish in which we placed 0.1 grams of NaHS and 2 mL of ultrapure water. Once we sealed the chamber, we gently shook it to put the crystal into contact with the water so that the gaseous H_2_S could be released into the chamber (Fig 6).

**Fig 6.**
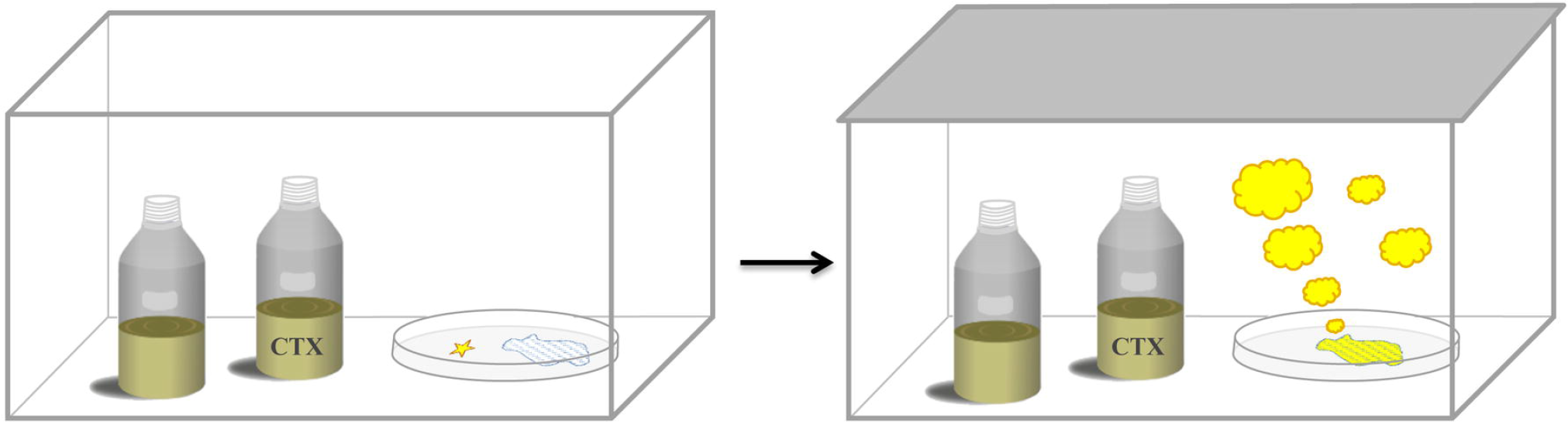
Method for incubation with chemical H_2_S. Samples were placed in a chamber, and next to it the NaHS stone (represented as a star) was placed on an empty Petri dish alongside 2 mL of ultrapure water (dashed blue area); after sealing the chamber, it is gently shaken to release the H_2_S to the atmosphere upon contact with H_2_O (B). CTX: cefotaxime.

With this experiment, we proved that the resistance of *E*. *faecalis* to cephalosporins could be reversed by pure H_2_S. In addition, we also showed that other *E*. *faecalis* strains are affected in the same way. Most notably, V583, the first vancomycin resistant clinical isolate, displayed the same phenotype (Fig 7).

**Fig 7.**
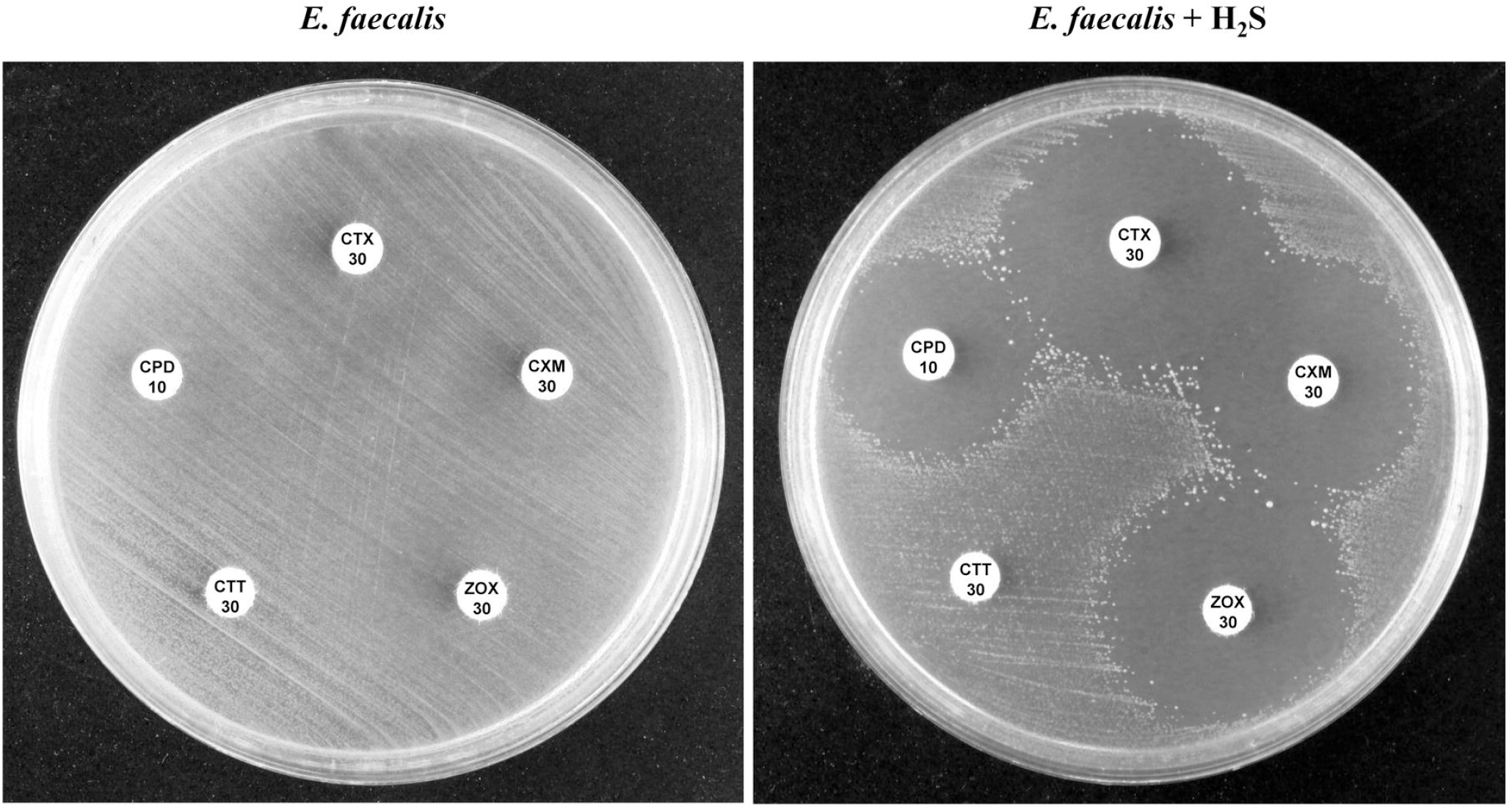
Antibiograms of *E*. *faecalis* in the presence of H_2_S. *E*. *faecalis* V583 was incubated in a chamber in the presence and abscence of NaHS 0.1 grams, for the antibiotics cefotaxime (CTX), cefuroxime (CXM), ceftizoxime (ZOX), cefotetan (CTT) and cefpodoxime (CPD).

### *E*. *faecium* cephalosporin resistance is also reverted in the presence of H_2_S

Next, we were interested in testing the effects of H_2_S on the other pathogenic enterococcus, *E*. *faecium*, as in the last decade this species has acquired even higher clinical relevance than *E*. *faecalis*, partially because most of the enterococcal vancomycin resistant clones currently detected are *E*. *faecium* [26]. We have verified that this species is also susceptible to the synergic effect of H_2_S with cephalosporins (Fig 8), further suggesting the potential application of this combination in clinical settings.

**Fig 8.**
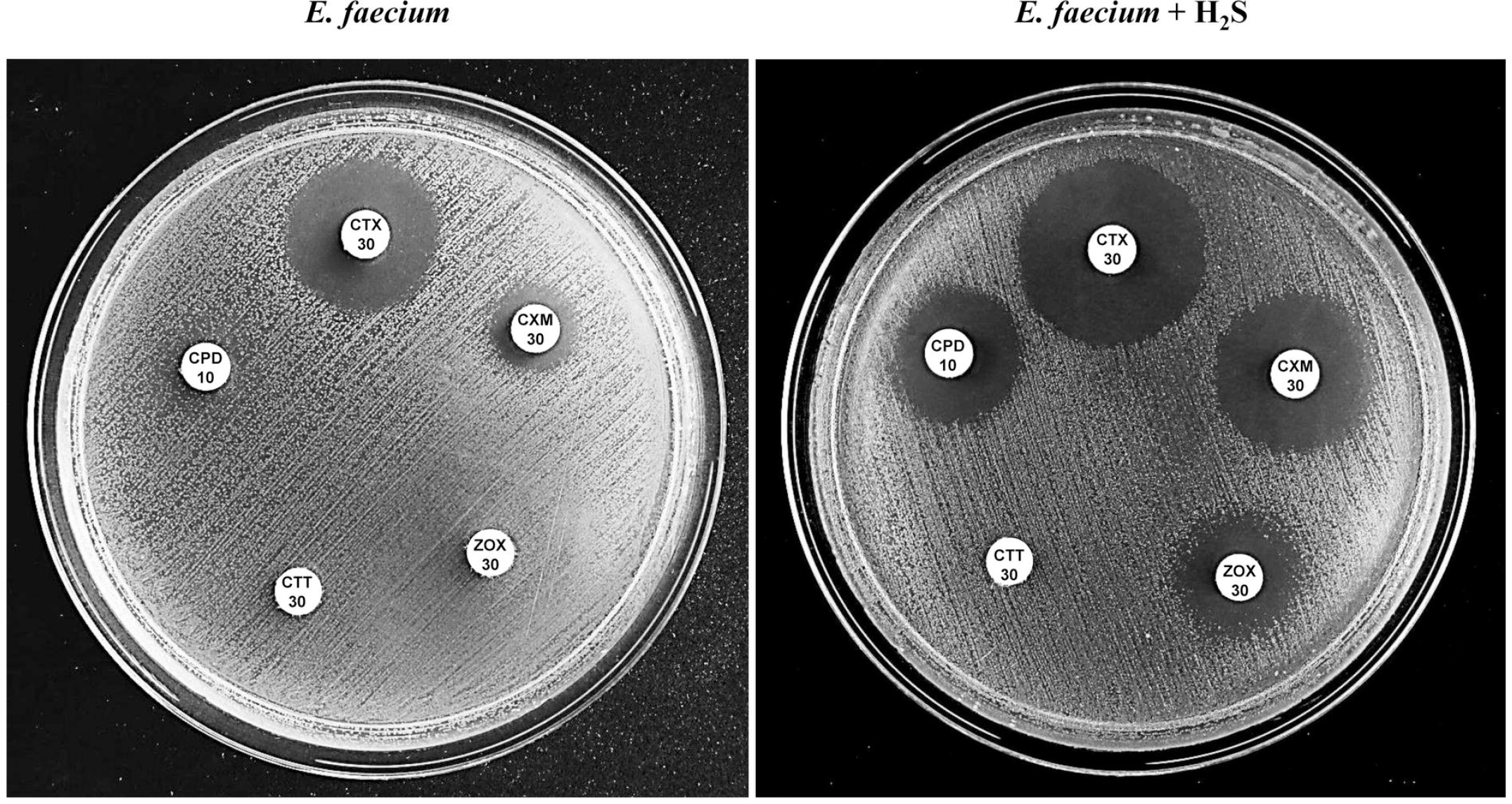
Antibiograms of *E*. *faecium* in the presence of H_2_S. *E*. *faecium* ATCC19434 was incubated in a chamber in the absence (left) and the presence (right) of 0.1 grams of NaHS, for the antibiotics cefotaxime (CTX), cefuroxime (CXM), ceftizoxime (ZOX), cefotetan (CTT) and cefpodoxime (CPD).

### Reversion of the intrinsic resistance of enterococci in the presence of H_2_S takes place specifically with methoxy-imino cephalosporins

As we can notice in Figs 3, 7 and 8, and Table 1, synergy between H_2_S and cephalosporins does not take place with every compound of this family of antibiotics. By performing antibiograms of a set of 20 cephalosporins, including 1^st^ to 4^th^ generation cephalosporins, we have observed that in the presence of H_2_S *E*. *faecalis* demonstrates a significant susceptibility against cefotaxime, ceftriaxone, cefuroxime, cefpodoxime and ceftizoxime (Table 2). These drugs, frequently used in clinical settings, contain a common methoxy-imino group in their structure (Fig 9), which is not present in other cephalosporins within the family, suggesting that this motif might be a key in the synergy displayed between H_2_S and cephalosporins.

**Table 2.** **Antibiograms of *E*. *faecalis* V583 strain, in the presence and absence of NaHS 0.1 grams**

|  | TSA | H<sub>2</sub>S 0.1 g |
| --- | --- | --- |
| Cefaclor 30 | 10 | 13 |
| Cefadroxil 30 | 13 | 13 |
| Cephalexin 30 | 8 | 8 |
| Cefalotin 30 | 16 | 16 |
| Cefamandole 30 | 14 | 15 |
| Cefazolin 30 | 14 | 16 |
| Cefepime 30 | 13 | 15 |
| <u>Cefotaxime 30</u> | 6 | 21 |
| Cefotetan 30 | 6 | 6 |
| Cefoxitin 30 | 6 | 6 |
| Cefpirome 30 | 22 | 25 |
| <u>Cefpodoxime 10</u> | 6 | 20 |
| Cefprozil 30 | 21 | 23 |
| Cefradine 30 | 12 | 13 |
| Cefsulodin 30 | 6 | 6 |
| <u>Ceftizoxime 30</u> | 6 | 16 |
| <u>Ceftriaxone 30</u> | 6 | 16 |
| <u>Cefuroxime 30</u> | 6 | 20 |

Methoxy-imino cephalosporins are underlined.

**Fig 9.**
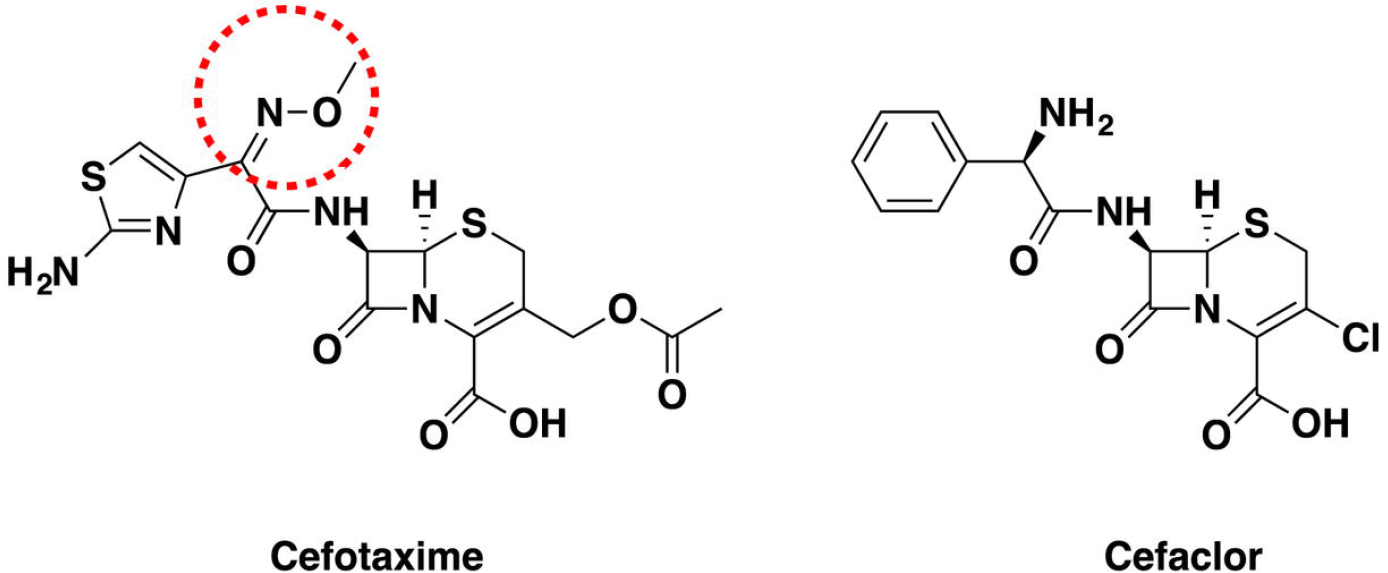
Chemical structure of cephalosporins. In cefotaxime the methoxy-imino residue is emphasized.

### H_2_S and cephalosporins kill enterococci

Finally, to further characterize the H_2_S-cephalosporins synergy, *E*. *faecalis* JH2-2 was grown in liquid media in the presence and absence of H_2_S and/or cefotaxime (CTX).

By performing a Two-way ANOVA (*α* = 0.05; *P* <0.001; *F* = 68,46; df = 17, 28) and a post-hoc Tukey–Kramer analysis, used for single-step multiple comparison of all pair of means, we observed that there is a strong synergistic effect between H_2_S and CTX (*P* < 0.001), not only by inhibiting bacterial growth in the presence of both compounds, but also by producing a bactericidal effect (Fig 10, S4 Table). With this analysis we confirmed that, in the concentration range used in our study, H_2_S does not affect bacterial growth on its own. Even if enterococci were resistant to cephalosporins, a minor effect of cefotaxime on bacterial growth was noticeable, but this is something that could be observed with many antibiotic discs in antibiograms or other antibiotic susceptibility tests.

**Fig 10.**
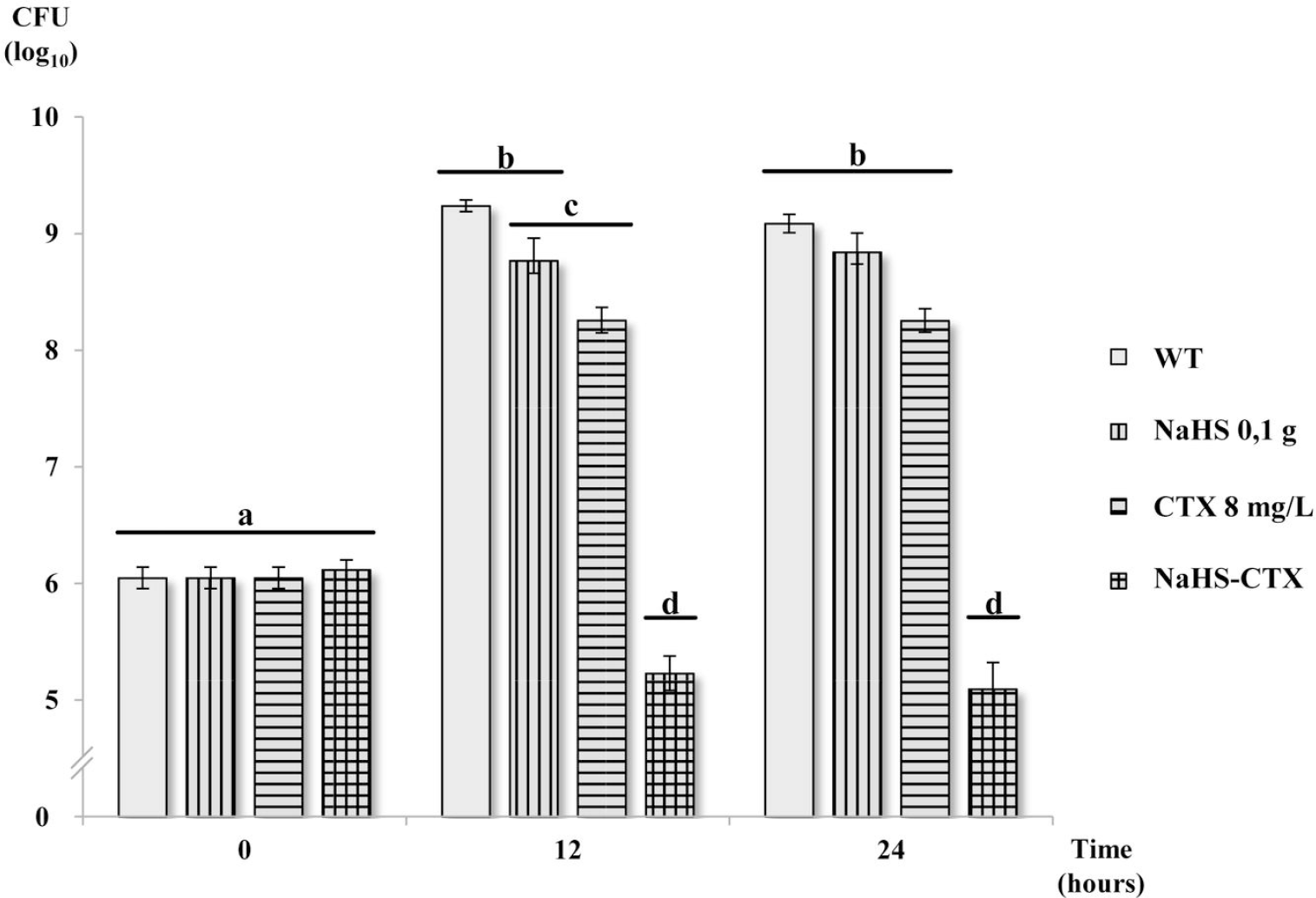
*E*. *faecalis* time killing curves. *E*. *faecalis* JH2-2 strain was incubated in TSB liquid media in the presence and absence of NaHS (in the atmosphere) and/or cefotaxime (CTX). Dilutions were plated at 0, 12 and 24 hours for Colony Forming Units (CFU) counting. Error bars represent the standard error of at least three independent experiments. Significant differences among samples are indicated as a, b, c, d. Samples with the same letter are not statistically different among them.

Preliminary experiments with cefaclor, a non-methoxy-imino cephalosporin, or with nalidixic acid, a quinolone against which *E*. *faecalis* is resistant, showed no synergy with H_2_S (data not shown), indicating that the displayed phenotype is specific of certain antibiotics.

### The persulfidome analysis of *E*. *faecalis* reveals potential targets for H_2_S

We carried out random transposition mutagenesis of *Enterococcus* under cephalosporin selective pressure, constructing a high-density transposon mutant library. However, no mutants were obtained using this approach. This suggests that a single gene alteration is not enough to prevent resistance reversion, or that this alteration has to take place in an essential gene. Therefore we wanted to address posttranslational changes of the protein that are induced by H_2_S.

Signaling by H_2_S is now widely linked to an oxidative posttranslational modification of cysteine residues (RSH) called persulfidation (RSSH) (alternatively S-sulfhydration) [27–29]. Exposure of cells to H_2_S results in an increase of protein persulfidation [27,30]. We performed biotin-tagging assay (BTA) to extract total persulfidated proteins [31,32]. As shown in Fig 11, an increase of protein persulfidation could be observed in bacteria treated with H_2_S.

**Fig 11.**
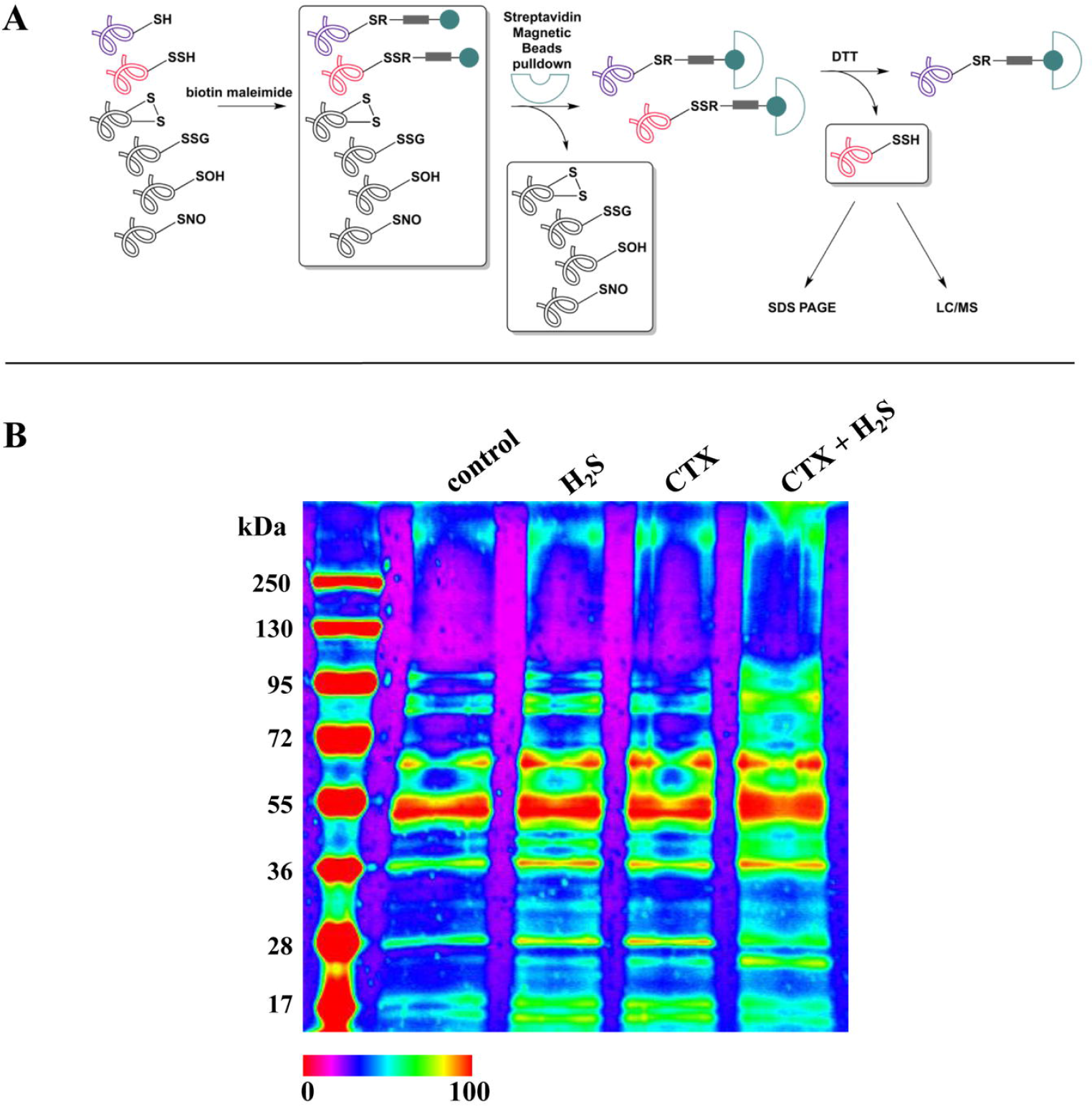
Identification of he complete persulfidome of *E. faecelis*. (A) Workflow used for the specific labeling of persulfide residues. Biotin maleimide binds only to thiol and persulfide motifs. Bound proteins are separated from others by streptavidin magnetic beads. DTT treatment specifically cleaves disulfide bridges with biotin maleimide, rendering the original S-sulfhydrated proteins available for subsequent procedures. (B) SDS-PAGE of *E*. *faecalis* total protein extraction incubated in the presence and absence of H_2_S and CTX.

The proteins were subjected to electrophoretic separation, followed by trypsin digestion and LC/MS/MS analysis. 66 cysteine-containing proteins were identified as persulfidated (Table 3, S7 Table). Among them, peptide ABC transporters, as well as proteins of unknown function are potential targets of H_2_S. Despite not being quantitative, to best of our knowledge, this is the first report of a persulfidome in bacteria. Further studies will be needed to determine the protein, or combination of proteins, that are responsible for the susceptibilization of enterococci to cephalosporins when exposed to H_2_S.

**Table 3.** Persulfidome of *E*. *faecalis*. List of proteins of *E*. *faecalis* JH2-2 detected to interact with H_2_S.

| Accession | Gene | Description |
| --- | --- | --- |
| Q833I9 | EF1963 | Phosphoglycerate kinase |
| Q839G8 | EF0201 | Elongation factor Tu |
| Q834N1 | EF1613 | Formate acetyltransferase |
| Q836Q8 | EF1050 | DNA-binding response regulator |
| Q833I8 | EF1964 | Glyceraldehyde-3-phosphate dehydrogenase |
| H7C7A0 | EF2739 | Alkyl hydroperoxide reductase, C subunit |
| Q837D6 | EF0907 | Peptide ABC transporter, peptide-binding protein |
| Q82ZF4 | EF3106 | Peptide ABC transporter, peptide-binding protein |
| P23530 | EF0710 | Phosphoenolpyruvate-protein phosphotransferase |
| Q839G9 | EF0200 | Elongation factor G |
| Q833J0 | EF1962 | Triosephosphate isomerase |
| Q831U9 | EF2398 | 30S ribosomal protein S2 |
| Q836C4 | EF1191 | DegV family protein |
| Q839C1 | EF0255 | L-lactate dehydrogenase 1 |
| Q836R2 | EF1046 | Pyruvate kinase |
| Q836Z4 | EF0949 | Phosphotransacetylase |
| Q837P9 | EF0784 | S-adenosylmethionine synthase |
| Q830Z3 | EF2621 | Uncharacterized protein |
| Q834N3 | EF1611 | Probable manganese-dependent inorganic pyrophosphatase |
| Q834M9 | EF1616 | CoA-binding domain protein |
| Q839E0 | EF0232 | 30S ribosomal protein S11 |
| Q838R7 | EF0368 | Aspartokinase |
| Q836Q9 | EF0149 | 6-phosphogluconate dehydrogenase, decarboxylating |
| Q839G1 | EF0209 | 50S ribosomal protein L2 |
| Q836D3 | EF1182 | S-ribosylhomocysteine lyase |
| Q831S5 | EF2425 | Phosphoglucomutase/phosphomannomutase family protein |
| Q834S3 | EF1560 | Uncharacterized protein |
| Q839H1 | EF0198 | 30S ribosomal protein S12 |
| Q837F1 | EF0891 | Aminotransferase |
| Q830Z2 | EF2622 | Uncharacterized protein |
| Q82ZJ2 | EF3064 | Polyribonucleotide nucleotidyltransferase |
| Q82Z22 | EF3257 | Oxidoreductase, pyridine nucleotide-disulfide family |
| Q838N8 | EF0401 | Pyrrolidone-carboxylate peptidase |
| Q835M1 | EF1356 | Dihydrolipoyl dehydrogenase |
| Q834H1 | EF1681 | Peptide methionine sulfoxide reductase MsrA |
| Q831A3 | EF2610 | ATP synthase subunit alpha |
| Q839I7 | EF0178 | ABC transporter, ATP-binding protein |
| Q835V2 | EF1270 | Ribosome maturation factor RimP |
| Q830B1 | EF2882 | Malonyl CoA-acyl carrier protein transacylase |
| Q834A7 | EF1763 | Protein translocase subunit SecA |
| Q830Y0 | EF2638 | Redox-sensing transcriptional repressor Rex 1 |
| Q839I2 | EF0185 | Phosphopentomutase |
| P37710 | EF0799 | Autolysin |

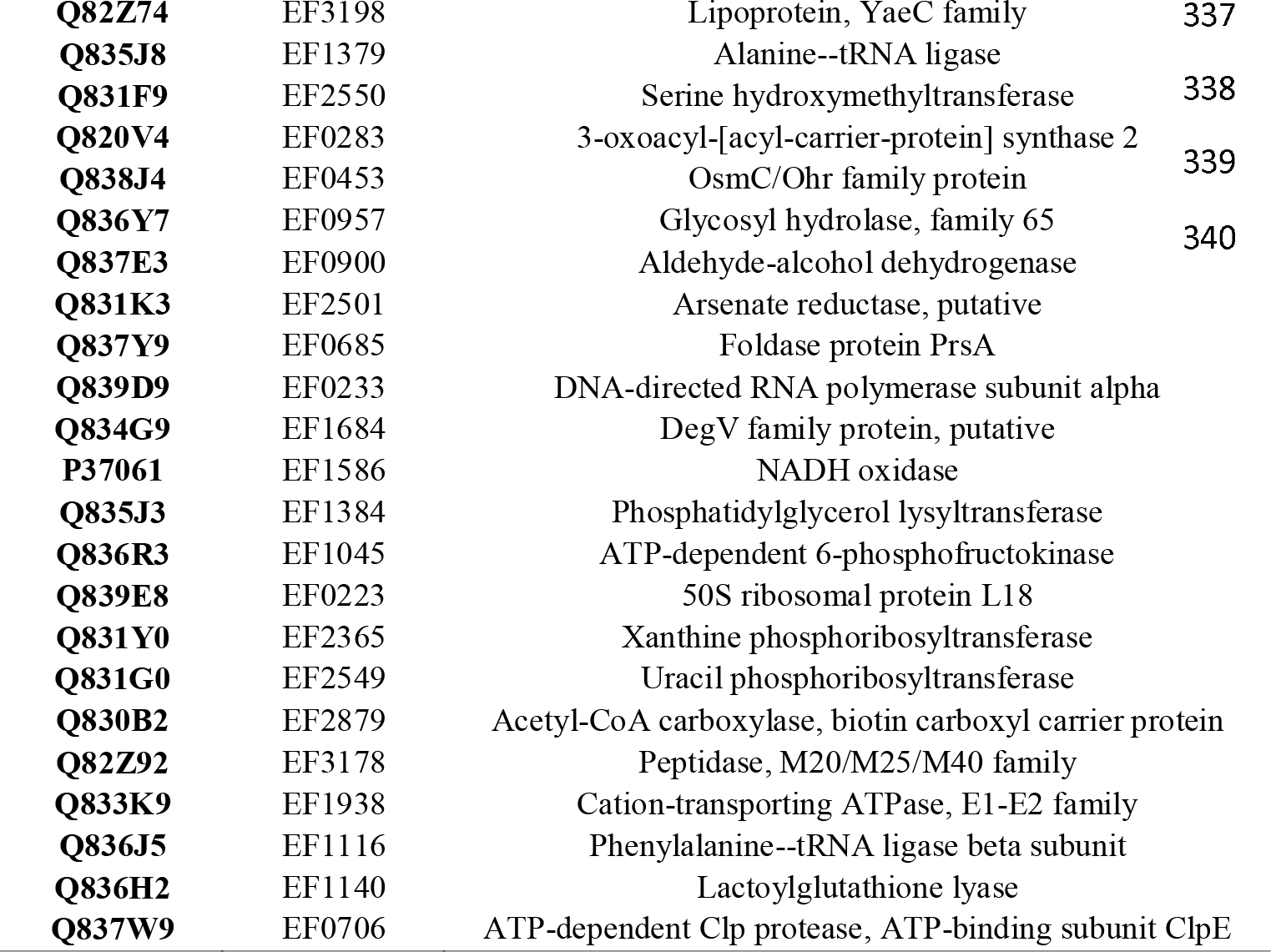

## Discussion

In this work we present significant progress in the study of H_2_S as a gasotransmitter and its link to antibiotic resistance.

*S*. Typhimurium is one of the classic H_2_S producing bacterial species, via its *phsABC* operon [18]. It is known that its capacity to reduce sulfur compounds provides *Salmonella* with a growth advantage when colonizing the digestive tract [17,33]. However, the physiology and involvement of the *phsABC* operon in antibiotic resistance, which has both a significant scientific and clinical interest, is not described.

The aim of this study was to understand the role and co-regulation of the H_2_S-producing pathways in *S*. Typhimurium, its possible role in microbial ecology and its effects on other bacteria. By knocking-out the *phsABC* operon, we have found that the PHS pathway is the predominant mechanism for H_2_S production in this species. Other pathways may or may not be active, but the lack of the PHS pathway is not compensated by other mechanisms.

We have demonstrated that the *phsABC* operon is not involved in own antibiotic resistance, even though it is the main source of H_2_S in *S*. Typhimurium. This suggests that the modifications in antibiotic resistance previously described in this species [20,34] are not due to the H_2_S itself. In fact, these authors state that the accumulation of H_2_S in their experiments is due to either a decrease in cysteine synthesis or an increase in its catabolism. Cysteine is a well-known inducer of the Fenton reaction and the H_2_S-mediated augmentation of antibiotic resistance is associated with oxidative stress protection [15]. This means that H_2_S itself may be irrelevant for antibiotic resistance in *S*. Typhimurium, which explains the absence of differences between WT and *Δphs* strains regarding every aspect we have tested (except for the production of H_2_S itself).

As we could not detect any intracellular role for H_2_S in *S*. Typhimurium, but we had verified that the *phsABC* operon was used to generate large amounts of H_2_S, we hypothesized that this H_2_S may have an extracellular effect, that is, on neighboring bacteria. Moreover, it is known that thiosulfate is abundant in the digestive tract [17], suggesting that the *phsABC* operon should be active.

When *E*. *faecalis* was tested, instead of observing an enhancement of antibiotic resistance by H_2_S, as previously described [15,20], we observed that this gas is capable of disrupting the intrinsic antibiotic resistance of *Enterococcus*. The MIC of cefotaxime in *E*. *faecalis* was dramatically decreased and resistance to other cephalosporins was similarly reverted, both in *E*. *faecalis* and *E*. *faecium* multiresistant enterococci, responsible for endocarditis and septicemia in hospitals worldwide, are ranked by the WHO among the pathogens for which effective therapies are critically needed [35]. In addition, the use of certain antibiotics is a predisposing factor for suffering an overgrowth and subsequent infections by *Enterococcus* [36,37]. Therefore, as pharmacological donors of H_2_S have already been developed [38–41], our work gives new hope for the treatment of enterococcal infections. H_2_S donor drugs could be administered together with cephalosporins to effectively kill multidrug resistant enterococci in patients. In addition, unraveling the mechanism of action of H_2_S inside *Enterococcus* could lead to the design of molecules that emulate H_2_S function.

In any case, considering that the supply of new antibiotics to the global market is virtually non-existent these days, the possibility of employing classical antibiotics that, until now, were unavoidably discarded beforehand to treat multiresistant infections, turns out encouraging in the fight against antibiotic resistance.

In our experiments, we observed that H_2_S exhibits bactericidal synergy with a select group of cephalosporins containing a methoxy-imino group in its structure. This residue improves the drug’s stability to β-lactamases and improves the ability of the drug to cross the external membrane of gram-negative bacteria [42]. Clearly, these properties do not account for its efficacy in the presence of H_2_S, as β-lactamase production by *Enterococcus* is anecdotic [43]. But, interestingly, many of the studies regarding cephalosporin resistance in enterococci were obtained with this specific group of methoxy-imino cephalosporins [44–49]. Thus, H_2_S might be affecting some of the pathways already described for *Enterococcus* cephalosporin resistance.

These pathways include a penicillin-binding protein that presents low affinity to these drugs [50], the MurAA enzyme involved in peptidoglycan synthesis [47], mutations of the β-subunit of the RNA polymerase [49], CroR/CroS and IreK/IreP two-component systems [45,48], or alterations in the thymidylate synthesis pathway [44], among others. Still, most of these studies do not unravel the final mechanisms governing cephalosporin resistance and they suggest a common currently undescribed mechanism [47,48,51]. Besides, in our experiments we have also observed an induction of aminoglycosides susceptibility by H_2_S (S3 Table), which it has not been observed by other authors. Therefore, H_2_S could be acting through a different pathway or upstream of a common mechanism, a theory that would be also supported by the fact that no known protein involved in cephalosporin resistance has been detected in the persulfidome (Table 3). In any case, aminoglycosides constitute another group of last resort antibiotics widely used in clinical settings and for which enterococci present a modest level of intrinsic resistance [52]. It would be interesting to study in depth this additional synergy between H_2_S and aminoglycosides, although it has not been object of study in this work, since the reversion of cephalosporin resistance in enterococci is more relevant from a clinical and microbiological perspective.

From an ecological perspective, it is noteworthy that, in raw meat, enterococci have been shown to successfully inhibit H_2_S production by sulfide-producing bacteria, presumably through enterocins or other biologically active compounds present in the supernatant [21], although no further studies have addressed this phenomenon. We have observed that, when exposed to H_2_S, *Enterococcus* does not acquire additional protection against antibiotics, but, in contrast, it is killed by a powerful group of antimicrobial agents, to which this species demonstrates a robust resistance. Therefore, this alteration of antibiotic resistance can explain why enterococci gain a selective advantage by blocking the H_2_S production of its neighbors. Enterococci constitute a small proportion of the physiological microbiota of both humans and animals [53]. Thus, both *Salmonella* and *Enterococcus* are pathogens which pursue the same niche colonization as part of their pathogenesis [17,37], which explains why both species would benefit if the other one is ousted. In addition, we have demonstrated how other H_2_S producing species, including *Proteus vulgaris* (which also share the same ecological niche), can also be involved in this interplay between bacteria.

Thus, this study highlights a new role for bacterial H_2_S production. We show how *Salmonella* and *Proteus*, through their H_2_S production, can obtain an unexpected advantage for niche competition (Fig 12). Aerial interaction among bacteria has been previously revealed, but, to the best of our knowledge, this is the first report about H_2_S aerial communication, showing how a bacterium can exert an effect with lethal consequences on another bacterium through H_2_S, a gasotransmitter with vital functions and abundant in the human body and to date considered a beneficial gas for bacteria.

**Fig 12.**
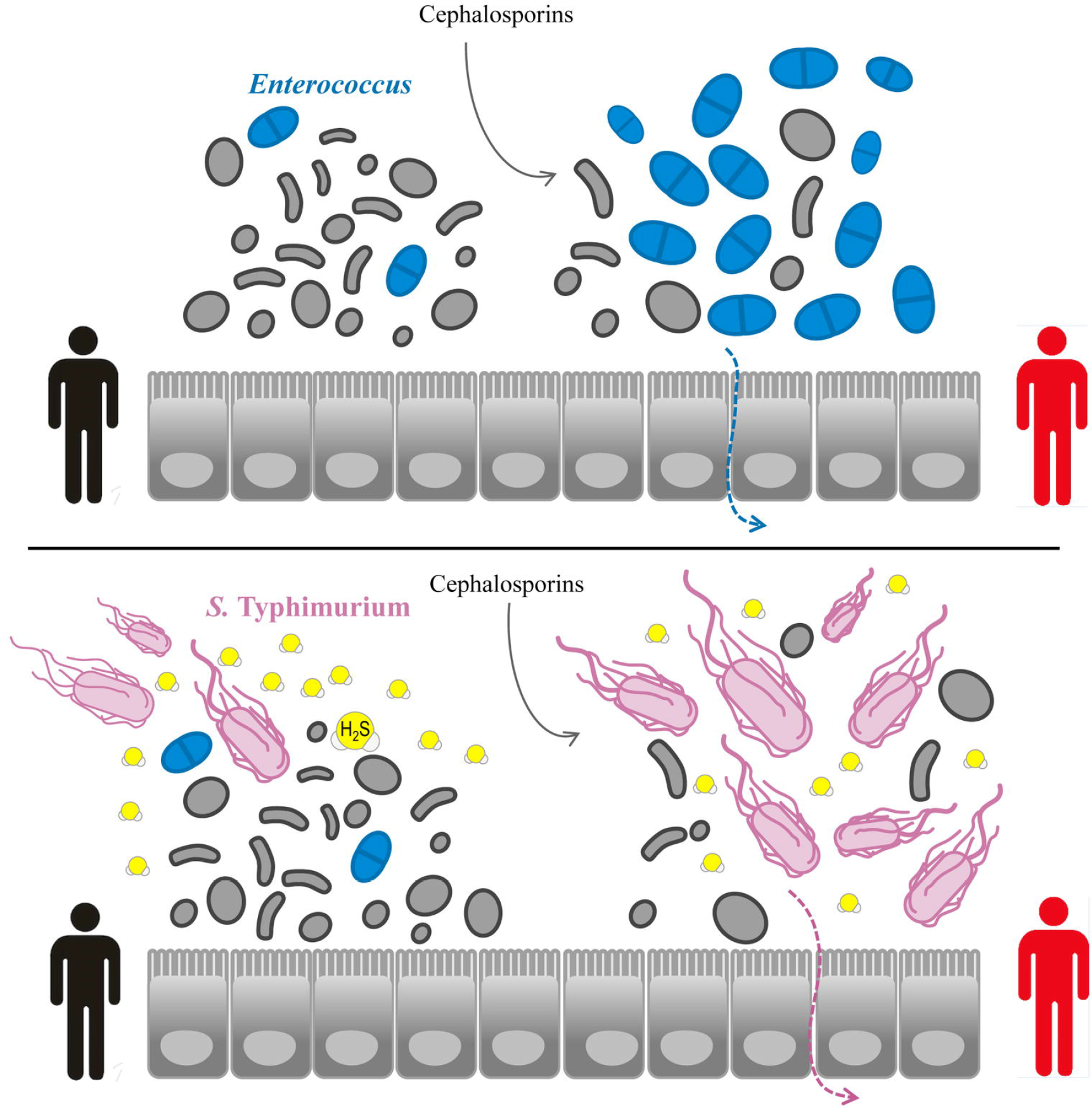
Niche competition of *Enterococcus* and H_2_S-producing bacteria. The use of cephalosporins or other antibiotics to which *E*. *faecalis* and *E*. *faecium* are resistant promotes these species overgrowth in the host’s digestive tract, facilitating their dissemination and the development of enterococcal infections (upper panel). Nevertheless, a cephalosporin-resistant *S*. Typhimurium strain in the niche could induce the killing of enterococci by H_2_S production, displacing the latter and colonizing the gut as a previous step for infection (bottom panel).

## Acknowledgements

We thank A. Hoefer for careful reading of the manuscript and G. Moyano for his assistance in statistics. We thank Eugeny Nudler for providing us with the *E*. *coli* MG1655 strain.

## Competing interests

The authors claim no conflict of interest.

## Funding

The funders had no role in study design, data collection and analysis, decision to publish, or preparation of the manuscript.

DTL and LC work have been supported by a FPU PhD scholarship from the Ministry of Education of Spain (AP2012-2130 and AP2008-00586, respectively).

SM was funded by a FEMS long-term fellowship.

NM work is funded by the EvoTAR and EFFORT European projects.

European Commission (EC) EFFORT-613754-FP7 Bruno Gonzalez-Zorn

European Commission (EC) EVoTAR-282004-FP7 Bruno Gonzalez-Zorn

MRF acknowledges the support from ATIP-Avenir and “Investments for the future” Programme IdEx Bordeaux, and the reference no. “ANR-10-IDEX-03-02”

## Authorship

Conceptualization: DTL, LC, BGZ. Investigation: DTL, NM, SM. Methodology: DTL, SM, SC, BGZ, MF. Project administration: BGZ. Supervision: BGZ, MF. Writing and review: DTL, BGZ, LC, MF.

## Competing interests

The corresponding author declares, on behalf of all authors, that there are no financial, personal or professional interests that could be construed to have influenced the work.

## Materials and Methods

### Strains, media and reagents

All experiments were performed in Tryptone-Soy Agar or Broth (TSA or TSB, respectively) purchased from Oxoid (Oxoid Ltd., UK), unless otherwise stated. Antibiotic disks were acquired from bioMérieux (bioMérieux SA, Marcy l’Etoile, France) and Oxoid. Antibiotic powder, sodium thiosulfate and other reagents were purchased from Sigma (Sigma-Aldrich Química SA, Spain). NaHS was acquired from Cayman (Cayman Chemical, USA).

The strains used in these experiments are listed in Table 4.

**Table 4.**
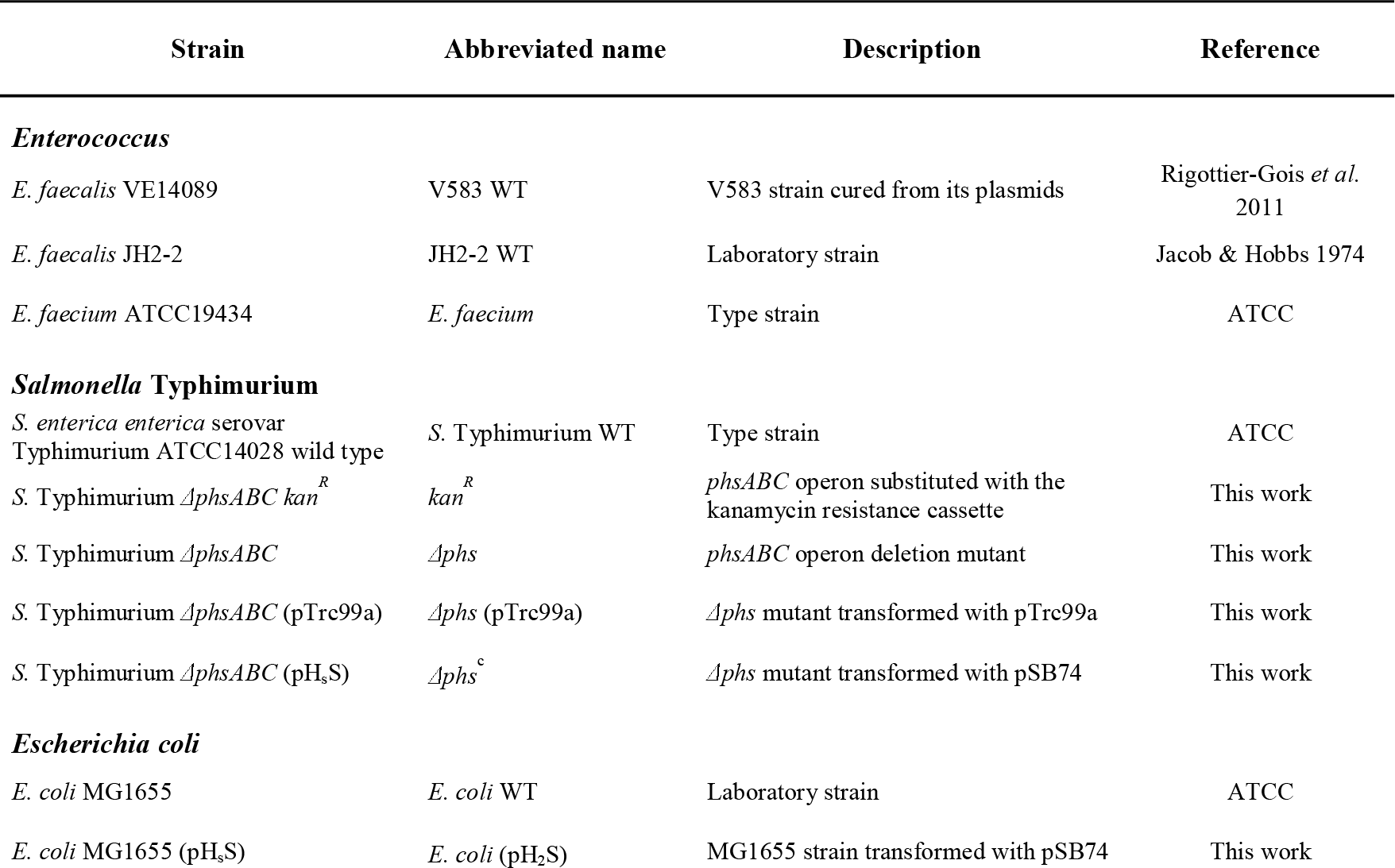
**Bacteria used in this study.**

Other strains
|  |  |  |  |
| --- | --- | --- | --- |
| *Bacillus cereus* CECT193 | *B. cereus* | Type strain | CECT |
| *Citrobacter freundii* | *C. freundii* | Strain I9 | Our laboratory |
| *Enterobacter cloacae* | *E. cloacae* | Strain RyC: L15 | Our laboratory |
| *Klebsiella pneumoniae* ATCC10031 | *K. pneumoniae* | Type strain | ATCC |
| *Listeria ivanovii* ATCC19119 | *L. ivanovii* | Type strain | ATCC |
| *Proteus vulgaris* CECT174 | *P. vulgaris* | Type strain | CECT |
| *Pseudomonas aeruginosa* | *P. aeruginosa* | Strain A26 | Our laboratory |
| *Staphylococcus aureus* COL | MRSA | Methicillin resistant *S. aureus* | Archer *et al.* 1994 |

**ATCC: American Type Culture Collection; CECT: Spanish Type Culture Collection.**

### Primers used and DNA manipulation

Reactives used for PCR were purchased from Biotools (Biotools, Madrid, Spain). TAQ polymerase (Biotools) or Phusion polymerase (Thermo Fisher Scientific Inc, USA) was used. PCR products were purified with the PCR purification kit (Qiagen, In.c, Chatsworth, CA). DNA was sequenced by Sanger (Secugen SL., Madrid, Spain). Plasmids were extracted with the QIAprep Miniprep and Midiprep kits.

Primers used are listed in S5 Table. All primers were purchased from Sigma-Aldrich.

### Plasmids used in this study

Plasmid pTrc99a was kindly donated by Professor Javier Turnay from the Biochemistry Department of the Complutense University of Madrid. pSB74 [23] was acquired from Addgene (plasmid #19591). Plasmid pKD13 (GenBank Accession number AY048744) was used as a PCR template in order to construct the deletion mutants and it contains an FLP recombination target (FRT)-flanked kanamycin resistance (*kan*) gene. Arabinose-induced Red helper plasmid pKD46 (GenBank Accession number AY048746) was used for the homologous recombination of the previously generated PCR products with the chromosome of *S*. Typhimurium ATCC14028. pCP20 FLP helper plasmid [54] was used for the elimination of the FRT-flanked resistance gene.

Plasmids used are listed in S6 Table.

### Construction of *S.* Typhimurium mutants

The *S*. Typhimurium *phsABC* deletion mutant was constructed as previously described [55,56]. Briefly, a PCR was performed using pKD13 DNA as the for the amplification of the FRT flanked kanamycin resistance (*kan*) gene subsequently used for mutant selection (3). The 5’-terminal deletion primer had a 50-nucleotide (nt) homologous extension that includes the *phsA* initiation codon, and the 20-nt 5’-ATTCCGGGGATCCGTCGACC-3’ priming site from pKD13. The 3’-terminal deletion primer consisted of a 50-nt homologous extension that includes 21 nt for the *phsC* C-terminal region, the termination codon and 29 nt downstream, and the 20-nt 5’-TGTAGGCTGGAGCTGCTTCG-3’ priming site from pKD13. The PCR product obtained, containing the FRT flanked kanamycin resistance gene in between *phsA* and *phsC* homologous regions, was transformed in *S*. Typhimurium ATCC14028 carrying the thermosensitive arabinose-induced plasmid pKD46. Upon addition of arabinose, kanamycin resistant colonies (that is, those who have had their *phsABC* operon replaced by the kanamycin cassette) were selected. pCP20 FLP helper plasmid was used for the elimination of the FRT-flanked resistance gene from the Δ*phsABC* deletion mutants. Correct in-frame deletion of the operon was checked with the primers phsABC_ext_F and phsABC_ext_R.

### Antibiograms and Minimal Inhibitory Concentration determination

Antibiograms were carried out in TSA media and interpreted following official guidelines (CLSI 2017; EUCAST 2017). Minimal Inhibitory Concentrations (MIC) of *E*. *faecalis* and *E*. *faecium* were determined by using E-test (bioMérieux) on TSA media. MICs of *S*. Typhimurium were determined in TSB by the broth microdilution method following official guidelines.

### Measurement of H_2_S production

Kligler media was purchased from bioMérieux. Tested strains freshly grown on TSA were inoculated on Kligler media and incubated overnight (o.n.), with the cap both tight and loose (to ensure aerobiosis conditions). Lead acetate paper strips were purchased from Sigma-Aldrich and used according to manufacturer’s instructions. Briefly, peptone water (purchased from Oxoid) tubes were inoculated with the tested strains. Then, between the cap and the inner wall of the tube we placed a strip, attached with adhesive tape and above the inoculated medium, and the cap was slightly tight. Samples were incubated for 18-24 hours at 37 °C.

### Confrontation experiments

For the confronting experiments (Fig 2), a distance of 1 cm between both bacteria was settled. To this end, 30 mL media plates were prepared, containing TSA, TSA supplemented with sodium thiosulfate (T-S) 2 or 20 mM, or TSA supplemented with cysteine (Cys) 2 mM. T-S and cysteine were prepared within the last seven days prior to the experiment. First, simulating the preparation of an antibiogram, a 0.5 McFarland turbidity suspension of the H_2_S producing species (or its mutant) was spread evenly over the entire surface of a TSA plate, supplemented with sodium thiosulfate T-S 2 mM, T-S 20 mM or Cys 2 mM prepared within the last seven days. Then, antibiograms of the strains to be tested were prepared in TSA, attending to official guidelines. Finally, lids were removed and both plates were fixed together with two pieces of adhesive tape, facing each other. The edges zone was “sealed” with air permeable Parafilm M (Bemis Company Inc, Oshkosh, WI). Plates were incubated at 37 °C for 24 hours maximum. Anaerobiosis conditions were achieved by using the GENBag system from bioMérieux.

### Growth curves

Growth curves of *S*. Typhimurium WT and its *Δphs* mutant were performed in Lennox broth media (Conda Laboratorios, Spain), as published [15]. Briefly, from an o.n. inoculum grown at 30 °C, a 1:100 dilution was performed in fresh media and samples were grown at 37 °C to 0.7-0.8 optical density at 600 nm (O.D.). Afterwards, a 1:100 dilution was carried out in 10 mL and samples were incubated at 37 °C and 150 rpm, measuring O.D. every hour. T-S 20 mM or different antibiotics were added when indicated.

### Experiments incubated with H_2_S

For experiments in the presence of H_2_S (Fig 6), *E*. *faecalis* antibiograms were prepared in TSA plates following official guidleline and were placed in a glass chamber of 2 liters volume. Next to them, we placed the base of an empty Petri dish in which we positioned 0.1 grams of NaHS and, separated, 2 mL of ultra-pure water. Then, we hermetically sealed the chamber and, only afterwards, NaHS and water were put in contact by gently shaking the chamber. Antibiograms were incubated for a maximum of 24 hours at 37 °C.

### Time killing experiments

10 μL from an o.n. culture of *E*. *faecalis* in TSB were inoculated to 15 mL glass bottles containing 10 mL of TSB. When required, antibiotics at their proper concentration were added. Bottles were incubated at 37 °C and 100 rpm. Those bottles that required H_2_S presence were placed in a glass hermetic chamber the same way as with experiments from the previous section (Fig 6). At 12 hours proper dilutions were plated for colony forming units (CFU) counting. From that moment on, all bottles were incubated under normal conditions until 24 hours, when dilutions were plated again.

### Random transposition mutagenesis

The library of mutants was obtained following the protocol designed by Zhang and collaborators [59]. Plasmid pZXL5 was transformed in the strain *E*. *faecalis* JH2-2. This thermosensitive plasmid contains: ColE and pWV01 origins of replication (for gram negative and gram positive, respectively), a *mariner* transposon (containing the *aax(6’)-apg(2’’)* gene for gentamicin resistance within), a nisin-inducible *mariner* transposase (with the system *nisK-nisR* and a chloramphenicol resistance cassette (*cat*).

Bacteria containing pZXL5 were incubated o.n. at 28 °C in BHI media containing gentamicin 300 mg/L and chloramphenicol 10 mg/L. Afterwards, a 1:2000 dilution was carried out in fresh BHI media containing 300 mg/L and nisin 0.025 mg/L to induce the random insertion of the transposon in the genome. Then, incubation of the samples at 150 rpm and 37 °C in the presence of gentamicin allows the specific selection of the bacteria possessing the gentamicin resistance gene (and, therefore, the transposon) in the chromosome.

Afterwards, a 1:2000 dilution in fresh media containing gentamicin was performed and bacteria were incubated at 37 °C and 150 rpm o.n., obtaining a library of mutants that was stored at −80 °C for future experiments.

By incubating in the presence of gentamicin, cefotaxime and H_2_S, we attempted to select only those mutants that resisted the combined action of H_2_S and cefotaxime, presumably because the transposon would have interrupted a pathway key in the susceptibilization of *Enterococci*.

### BTA method for persulfide detection

Bottles containing fresh TSB or TSB-CTX 8 μg/mL were inoculated with an o.n. inoculum of *E*. *faecalis* JH2-2, in a 1:1000 proportion. Samples were incubated for 5.5 hours at 37 °C and 100 rpm. Then, each sample was split in two and, to one of each set, H_2_S 250 μM (freshly prepared as previously described) was added. The four bottles were incubated for 30 minutes at 37 °C and 100 rpm. Then, samples were centrifuged for 3 minutes at 6000 rpm and supernatant was discarded. Samples were washed once with PBS, centrifuged and supernatant discarded again. Then, each sample was mixed with 1 mL of lysis buffer, containing HENP buffer (pH 7.41), SDS 1%, 10 μL of protease inhibitor 1% and 100 μM of biotin maleimide. To ensure complete lysis, samples were sonicated (3 cycles of 1 minute at 190 MHz and 2 minutes of pause). Then, samples were incubated for 60 minutes at 37 °C.

After incubation, 20 mM of freshly prepared N-ethylmaleimide (NEM) was added to block the unreacted cysteines and the samples were incubated o.n. at room T^a^ with rotating shaking. During all the procedure the lysis buffer and the samples were protected from light.

Proteins were precipitated with CHCl_3_/MeOH/H_2_O precipitation (1/4/4, v/v), washed twice and resuspended in HEPES buffer pH 7.4 containing SDS up to 0.5%. Samples were adjusted to equal protein concentration. Biotinylated proteins were bound to streptavidin agarose beads, and persulfidated targets eluted by DTT 0.5 mM (Fig 11).

Eluted proteins were either resolved by SDS electrophoresis and visualized with silver staining or subjected to in-gel trypsinization and subsequent LC/MS analysis. Experiments were performed in triplicates.

### nanoLC/MS

Protein samples were run on SDS-PAGE (10 %) but protein separation was stopped once proteins have entered the resolving gel. After colloidal blue staining, each lane was was cut into 1 mm × 1 mm gel pieces. The gel pieces were destained in 25 mM ammonium bicarbonate 50% ACN, rinsed twice in ultrapure water and shrunk in ACN for 10 min. After ACN removal, gel pieces were dried at room temperature, carbamidomethylated and covered with the trypsin solution (10 ng/μL in 50 mM NH_4_HCO_3_), rehydrated at 4°C for 10 min, and finally incubated overnight at 37°C. The gel pieces were then incubated for 15 min in 50 mM NH4HCO3 at room temperature with rotary shaking. The supernatant was collected, and an H_2_O/ACN/HCOOH (47.5:47.5:5) extraction solution was added onto the gel pieces for 15 min. The extraction step was repeated twice. The pooled supernatants were dried in a vacuum centrifuge and then resuspended in 40 μL of water acidified with 0.1% HCOOH. Peptide mixture was analyzed on an Ultimate 3000 nanoLC system (Dionex, Amsterdam, The Netherlands) coupled to an Electrospray Q-Exactive quadrupole Orbitrap benchtop mass spectrometer (Thermo Fisher Scientific, San Jose, CA). Ten microliters of peptide digests were loaded onto a 300-μm-inner diameter × 5-mm C18 PepMapTM trap column (LC Packings) at a flow rate of 30 μL/min. The peptides were eluted from the trap column onto an analytical 75-mm id × 15-cm C18 Pep-Map column (LC Packings) with a 4–40% linear gradient of solvent B in 108 min. Mobile phases were a mix of solvent A (0.1% formic acid in 5% ACN) and solvent B (0.1% formic acid in 80% ACN). The separation flow rate was set at 300 nL/min. The mass spectrometer operated in positive ion mode at a 1.9-kV needle voltage. Data were acquired using Xcalibur 2.2 software in a data-dependent mode. MS scans (m/z 300–2000) were recorded at a resolution of R = 70000 (@ m/z 200) and an automatic gain control (AGC) target of 3 × 106 ions collected within 100 ms. Dynamic exclusion was set to 30 s and the top 12 ions were selected from fragmentation in higher-energy collisional dissociation (HCD) mode. MS/MS scans with a target value of 1 × 105 ions were collected with a maximum fill time of 120 ms and a resolution of R = 35000. Additionally, only +2 and +3 charged ions were selected for fragmentation. Other settings were as follows: neither sheath nor auxiliary gas flow; heated capillary temperature, 270°C; normalized HCD collision energy of 27% and an isolation width of 2 m/z.

### Database search and results processing

Data were searched by SEQUEST through Proteome Discoverer 1.4 (Thermo Fisher Scientific Inc.) against a subset of the 2017-03 version of UniProt database restricted to
*Enterococcus faecalis* Reference Proteome Set (3240 entries). Spectra from peptides higher than 5000 Da or lower than 350 Da were rejected. The search parameters were as follows: mass accuracy of the monoisotopic peptide precursor and peptide fragments was set to 10 ppm and 0.02 Da respectively. Only b– and y-ions were considered for mass calculation. Oxidation of methionines (+16 Da) was considered as variable modification and carbamidomethylation of cysteines (+57 Da) as fixed modification. Two missed trypsin cleavages were allowed. Peptide validation was performed using Percolator algorithm [60] and only “high confidence” peptides were retained corresponding to a 1% False Positive Rate at peptide level.

### Statistical analysis

The statistical analysis for this paper was generated using SAS software Pre-production version 9.0 (SAS Institute Inc., Cary, NC, USA).

## Supplementary material

### S1 Fig. *S.* Typhimurium WT and *Δphs* growth curves

Experiments were carried out both in the presence and absence of thiosulfate (T-S) 20 mM (A) and in the presence of ampicillin 0.5 mg/L (B). The error bars represent the standard error of three independent experiments. Minimal Inhibitory Concentrations of different antibiotics (C). The results show the mean of three independent experiments.

**S1 Table.** **Antibiograms of *Salmonella* Typhimurium WT and Δ*phs* strain, in Tryptone Soy Agar (TSA), in the presence and absence of thiosulfate 20 mM (T-S).**

|  |  | ***S. Typhimurium*** |  | ***S. Typhimurium*** |  |
| --- | --- | --- | --- | --- | --- |
|  |  | **WT** |  | **$\Delta phs$** |  |
|  |  | TSA | TSA +<br>T-S 20 | TSA | TSA +<br>T-S 20 |
|  | **Amox. + Clavulanic acid 30** | 31 | 28 | 32 | 29 |
|  | **Cefaclor 30** | 31 | 30 | 29 | 31 |
|  | **Cefotaxime 30** | 34 | 34 | 34 | 34 |
|  | **Cefpodoxime 10** | 28 | 29 | 30 | 27 |
|  | **Cefoxitin 30** | 26 | 25 | 29 | 25 |
|  | **Imipenem 10** | 33 | 32 | 33 | 32 |
|  | **Aztreonam 30** | 38 | 34 | 34 | 34 |
|  | **Apramycin 15** | 13 | 10 | 13 | 9 |
|  | **Gentamicin 10** | 16 | 14 | 17 | 14 |
|  | **Neomycin 30** | 16 | 14 | 15 | 13 |
|  | **Netilmicin 30** | 19 | 17 | 19 | 17 |
|  | **Estreptomycin 10** | 12 | 7 | 12 | 7 |
|  | **Ciprofloxacin 10** | 35 | 37 | 35 | 36 |
|  | **Sulpha - Trimetoprim 25** | 29 | 30 | 29 | 30 |
|  | **Tetracycline 30** | 27 | 27 | 26 | 27 |
|  | **Clindamycin 2** | 6 | 6 | 6 | 6 |
|  | **Colistin 10** | 14 | 14 | 13 | 14 |
|  | **Cloramphenicol 50** | 25 | 27 | 28 | 28 |
|  | **Erythromycin 15** | 7 | 6 | 6 | 7 |
|  | **Mupirocin 5** | 6 | 6 | 6 | 6 |
|  | **Rifampin 5** | 9 | 7 | 9 | 9 |
|  | **Vancomycin 30** | 6 | 6 | 6 | 6 |
|  | **Fosfomycin 200** | 36 | 30 | 33 | 31 |

**S2 Table.**
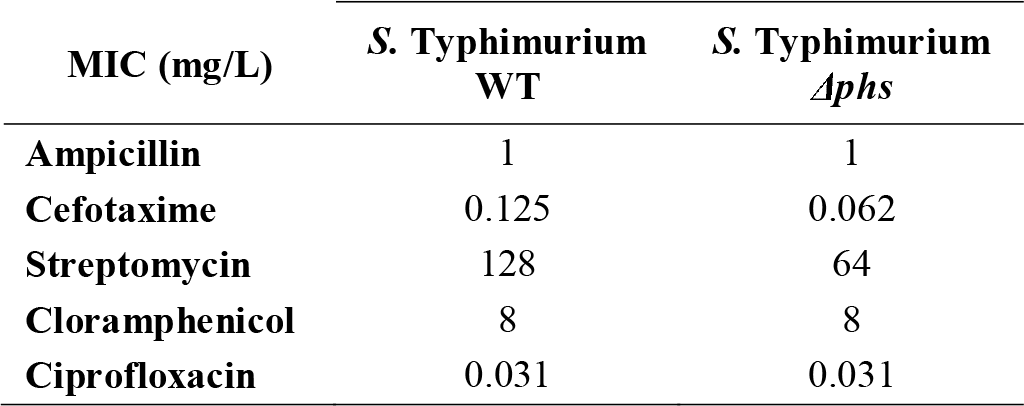
**Minimal Inhibitory Concentration (MIC) for different antibiotics in *S.* Typhimurium WT and its *Δphs* mutant.**

**S3 Table.**
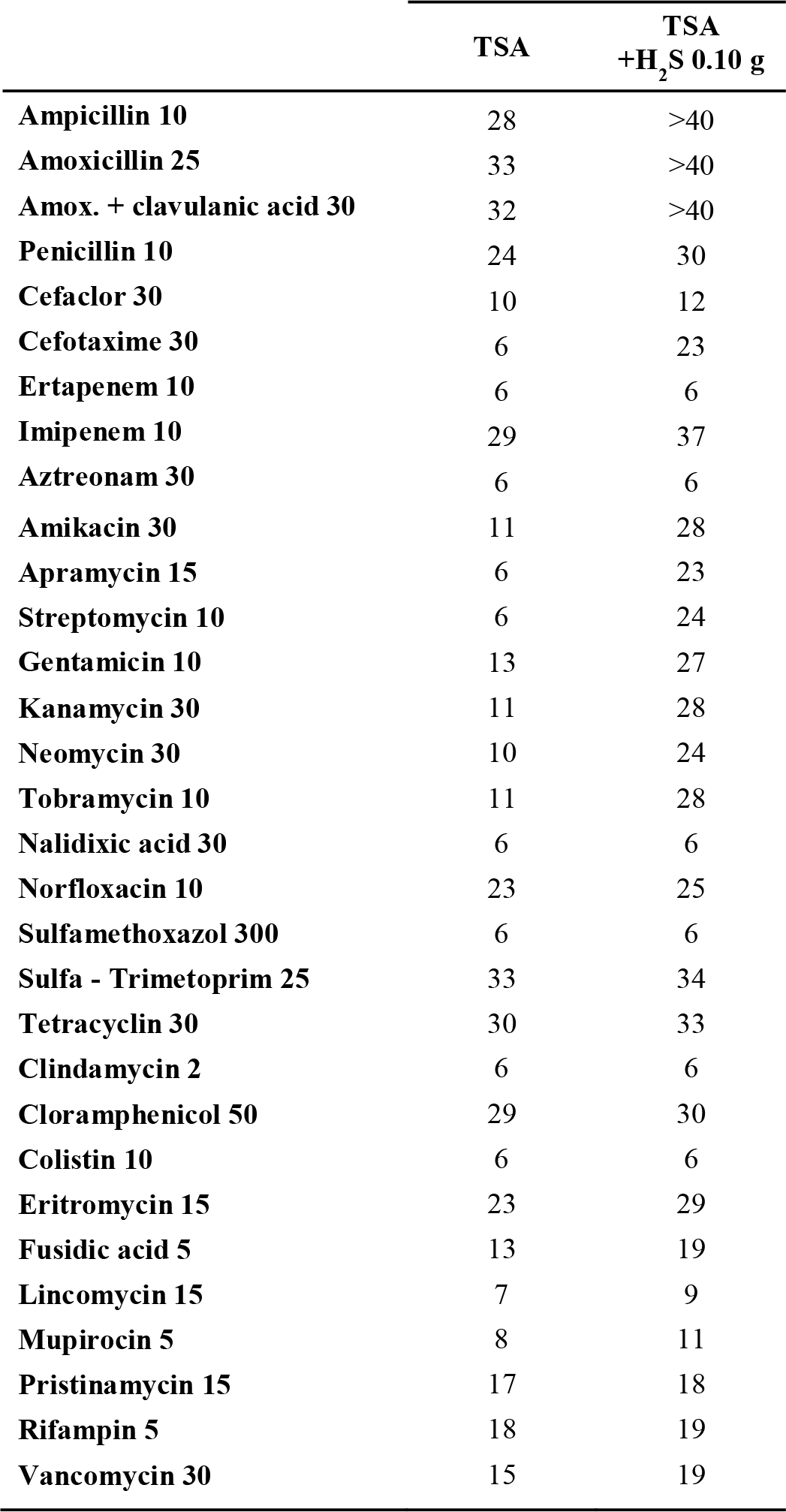
**Antibiograms of *E*. *faecalis* V583 strain, in Tryptone Soy Agar (TSA), in the presence and absence of NaHS 0.10 grams.**

**S4 Table.**
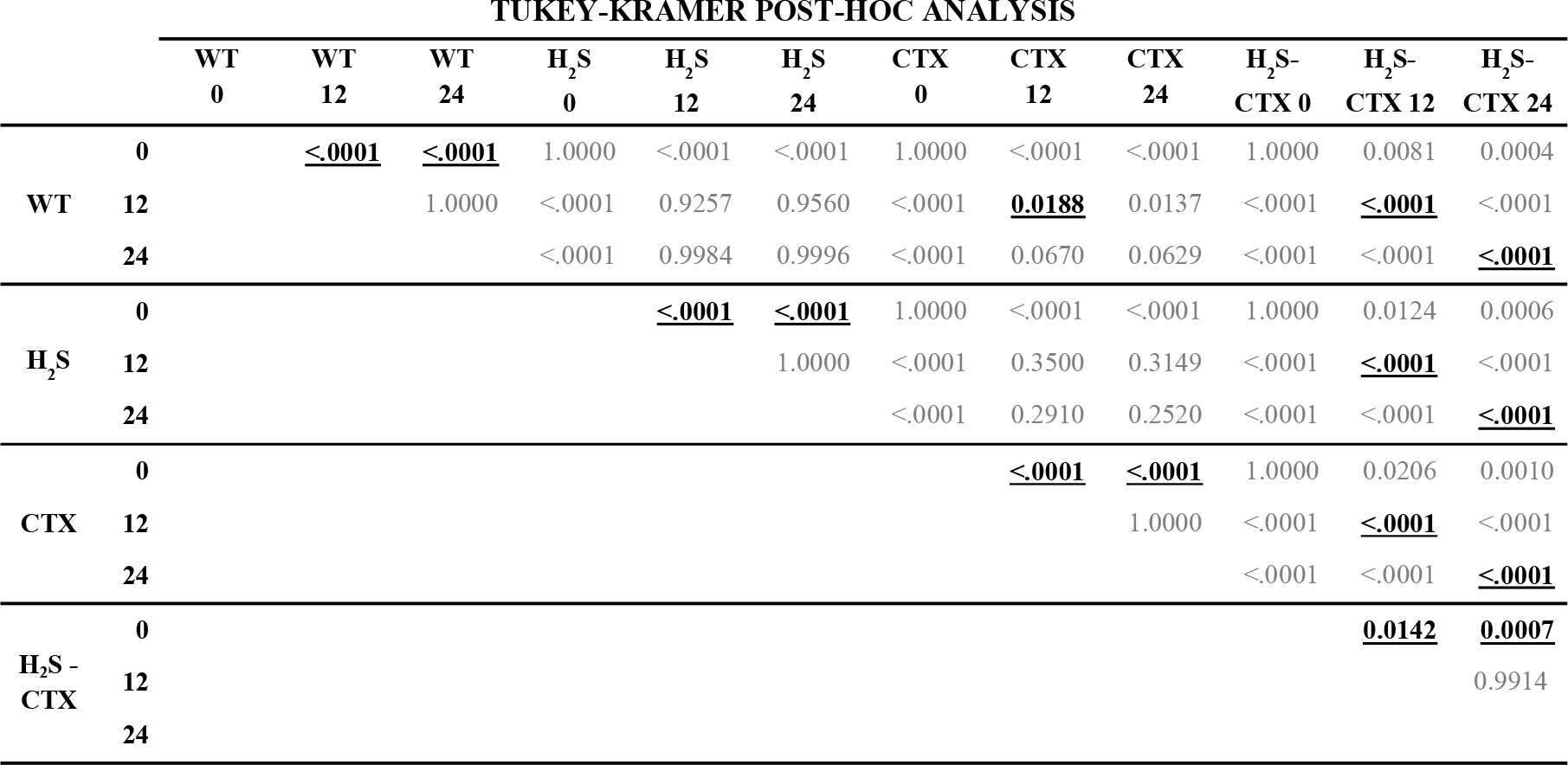
**Tukey-Kramer post-hoc analysis results. Statistically significant results of interest are highlighted.**

**S5 Table.**
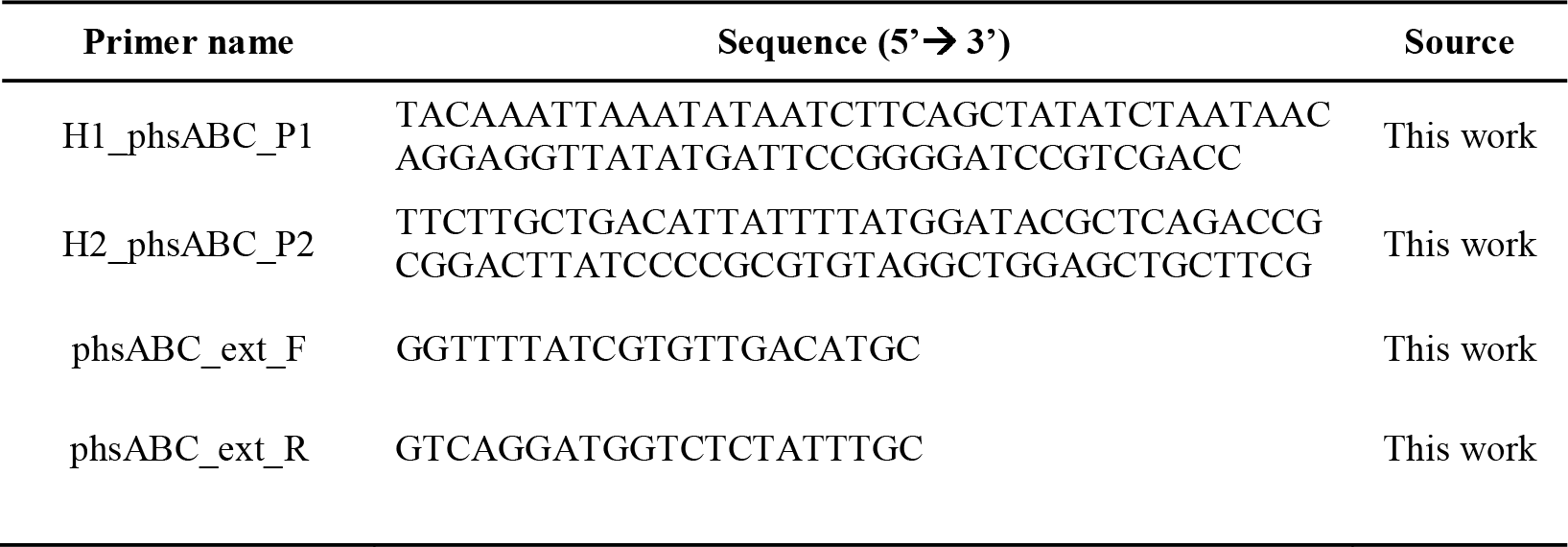
**Primers used in this study.**

**S6 Table.** **Plasmids used in this study.**

| Plasmid name | Description | GenBank Access number | Reference |
| --- | --- | --- | --- |
| <u>In-frame genes deletion</u> |  |  |  |
| pKD46 | Thermosensitive helper plasmid. It carries the $\lambda$ phage Red recombination system. Ara<sup>Ind</sup>, Amp<sup>R</sup> | AY048746 | Datsenko & Wanner 2000 |
| pKD13 | Kanamycin resistance cassette carrier plasmid. | AY048744 | Datsenko & Wanner 2000 |
| pCP20 | Thermosensitive helper plasmid. It carries the FLP recombinase. Amp<sup>R</sup> | - | Cherepanov & Wackernagel 1995 |
| <u>H<sub>2</sub>S production</u> |  |  |  |
| pTrc99a | Empty vector |  | Bang *et al.* 2000 |
| pSB74 (pH<sub>2</sub>S) | pTrc99a + *phsABC* | - | Bang *et al.* 2000 |
| <u>Random transposition mutagenesis</u> |  |  |  |
| pZXL5 | Thermosensitive plasmid. It carries a *mariner* transposon (Gm<sup>R</sup>) and a *mariner* transposase. Chlor<sup>R</sup> | JQ088279.1 | Zhang *et al.* 2012 |

